# Inter-organelle crosstalk supports acetyl-coenzyme A homeostasis and lipogenesis under metabolic stress

**DOI:** 10.1101/2022.09.24.509326

**Authors:** Ramya S Kuna, Avi Kumar, Hector Galvez, Karl A Wessendorf-Rodriguez, Courtney R Green, Grace H McGregor, Thekla Cordes, Reuben J Shaw, Robert U Svensson, Christian M Metallo

## Abstract

Proliferating cells rely on acetyl-CoA to support membrane biogenesis and acetylation. Several organelle-specific pathways are available for provision of acetyl-CoA as nutrient availability fluctuates, so understanding how cells maintain acetyl-CoA flux under such stresses is critically important. To this end we applied ^13^C isotope tracing cell lines deficient in these mitochondrial (ATP-citrate lyase; ACLY-), cytosolic, (acetyl-CoA synthetase (ACSS2-), and peroxisomal (peroxisomal biogenesis factor 5; PEX5-) dependent pathways. ACLY knockout in multiple cell lines reduced fatty acid synthesis and increased reliance on extracellular lipids or acetate. Knockout of both ACLY and ACSS2 (DKO) severely stunted but did not entirely block proliferation, suggesting alternate pathways can support acetyl-CoA homeostasis. Metabolic tracing and PEX5 knockout studies link peroxisomal oxidation of exogenous lipids as a major source of acetyl-CoA for lipogenesis and histone acetylation, highlighting a role for inter-organelle crosstalk in supporting cell survival in response to nutrient fluctuations.

**Teaser:** We quantify how acetyl-CoA metabolism is supported by distinct pathways spanning mitochondria, cytosol, and peroxisomes using comprehensive tracing applied to knockout cells.

## Introduction

Acetyl-CoA is a critical precursor for acetylation and lipid synthesis associated with membrane biogenesis, energy storage, and protein modification (*1*). Central carbon metabolism is wired with several redundancies in place to maintain acetyl-CoA homeostasis under conditions of stress, with mitochondria, peroxisomes, and cytosolic acetate activation all contributing to this metabolic pool. While both healthy tissues and tumors readily incorporate lipids from the diet (*1*), *de novo* lipogenesis (*DNL*) is required for some tumors and cell types to proliferate (*2*, *3*), how each of these pathways and organelles fuel lipogenesis is not fully clear. Fatty acids are synthesized from two carbon acyl-units provided by cytosolic acetyl-CoA, which is metabolized to malonyl-CoA by acetyl-CoA Carboxylase (ACC) and used by fatty acid synthase (FASN) to generate long-chain fatty acids. These fatty acids can be further elongated, desaturated, and incorporated into lipids within the endoplasmic reticulum (ER). In the context of cancer, FASN inhibition reduces body weight (*4*), oncogene expression (*5*), and *in vivo* growth (*6*) of human breast cancer cell lines in mice. Knockout and pharmacological inhibition of acetyl-CoA carboxylase also compromise tumor growth in xenograft and genetically-engineered mouse models of lung cancer while synergizing with chemotherapy (*7*).

ATP-citrate lyase (ACLY) is the major conduit through which mitochondrial metabolism supports acetyl-CoA generation, which is important for lipid synthesis as well as epigenetic regulation and DNA repair via histone acetylation (*8–10*). Upregulation of ACLY expression has been identified in a number of numerous cancers upregulate ACLY expression compared to surrounding healthy tissue, including breast, liver, bladder, stomach, colorectal, glioblastoma, and lung cancers (*11*–*17*), and in patients with gastric adenocarcinoma (*18*) and lung adenocarcinomas (*17*), overexpression and activation of ACLY is associated with poor prognosis. Further, both pharmacological inhibition and genetic knockdown of ACLY have been demonstrated to reduce lipogenesis and cancer cell proliferation (*19–26*), supporting the link between ACLY activity and tumor growth. On the other hand, nutrient availability and mitochondrial metabolism are sometimes compromised in cells within the tumor microenvironment, and pathways like reductive carboxylation are often unable to fully compensate (*27*, *28*). Therefore, characterization of alternative acetyl-CoA sources may useful for understanding cancer cell survival and resistance mechanisms.

The acetyl-CoA synthetase family of enzymes (ACSS1, ACSS2, and ACSS3) are the rate limiting enzymes involved in acetate activation to acetyl-CoA in mammalian cells (*29*). While ACSS1 and ACSS3 are mitochondrially expressed and tissue-restricted (in the case of ACSS3), ACSS2 is localized in the cytosol (*30*). ACSS2 has garnered interest as a cancer target as high expression of ACSS2 is associated with poor prognosis in breast, glioblastoma, ovarian, and lung cancers (*31*–*33*). Nutrient deprivation, including hypoxia and lipid depletion, upregulate ACSS2 expression and acetate catabolism for DNL in cancer cells, suggesting that ACSS2 provides a mode of metabolic plasticity to cancer cells under stress (*34*). Hepatocyte selective depletion of ACSS2 or ACLY decreased liver acetyl-CoA levels in obese mice (*35*). Furthermore, mouse embryonic fibroblasts (MEFs) deficient in ACLY upregulate ACSS2 protein expression and increase utilization of acetate for cytosolic acetyl-CoA generation (*36*), highlighting the essentiality of acetyl-CoA and metabolic flexibility of proliferative cells.

Peroxisomes generate acetyl-CoA via β-oxidation of fatty acids, most often very long-chain (VLCFA) or branched-chain fatty acids (BCFA), which contribute to malonyl-CoA and lipids in the heart and liver (*37–40*). Peroxisomal enzymes involved in oxidation of branched chain fatty acids (HSD17B4, ACOX3) are overexpressed in cancerous prostate tissue compared to paired healthy tissue (*41*), suggesting this organelle may support generation of acetyl-CoA in tumors. However, the extent to which peroxisomal β-oxidation of fatty acids serves as a source of lipogenic acetyl-CoA in cancer cells under nutrient stress warrants further exploration.

Here we apply mass spectrometry, stable isotope tracing and isotopomer spectral analysis (ISA) to analyze the metabolic rewiring that occurs when acetyl-CoA synthesis is systematically compromised in cancer cells. We genetically engineered cancer cell lines deficient in ACLY and/or ACSS2 using CRISPR/Cas9. While knockout of ACLY decreased *DNL* under basal conditions, cells are able to sustain FASN flux and/or growth through use of exogenous acetate or serum lipids. On the other hand, knockout of both ACLY and ACSS2 induces a growth limiting stress regardless of medium nutrient status. Finally, we observe that ACLY-deficient cells use peroxisomal β-oxidation as a significant source of lipogenic acetyl-CoA and are sensitive to peroxisomal biogenesis factor 5 (PEX5) knockout or pharmacological inhibition of peroxisomal β-oxidation. Collectively, our study highlights the rewiring of organelle-specific acetyl-CoA metabolism in fueling DNL when mitochondrial (ACLY), cytosolic (ACSS2), and peroxisomal (PEX5) CoA sources are compromised in cancer cells.

## Results

### ACLY deficiency compromises fatty acid synthesis and drives lipid-dependence of cell lines

*ACLY*-knockout (*ACLY*-KO) clones were generated using CRISPR/Cas9 in A549 human NSCLC cells, 634T murine NSCLC cells, HepG2 hepatocarcinoma cells, and HAP1 cells (Fig. 1B, S1A). We observed increased protein expression of ACSS2 in the A549 and HepG2 *ACLY*-KO cell lines (Fig. 1B, S1A), consistent with previous results in MEFs (*36*). The growth rate of the *ACLY*-KO A549, 634T, and HAP1 cells was marginally lower compared to wildtype cells, while HepG2 *ACLY*-KO cells showed an over 50% reduction in growth rate compared to the wildtype cells (Fig. 1C, S1B). In A549 cells, *ACLY*-KO resulted in decreased glucose uptake/lactate secretion and increased glutamine uptake/glutamate secretion, suggesting rewiring of central carbon metabolism (Fig. 1D). To better understand how ACLY-deficiency alters intracellular metabolism, we cultured A549 clones in the presence of [U-^13^C_6_]glucose or [U-^13^C_5_]glutamine and quantified isotope enrichment in downstream metabolite pools. Enrichment of TCA cycle intermediates from glutamine in A549 *ACLY*-KO cells was elevated, consistent with extracellular flux results (Fig.1E). These findings may indicate that ACLY-KO induces oxidative stress, as increased glutamine catabolism has been found to be integral to maintaining redox homeostasis in cancer cells through multiple mechanisms (*42*, *43*). On the other hand, glucose enrichment in malate and aspartate (a readout of oxaloacetate) was increased, likely due to decreased generation of acetyl-CoA via ACLY and glutamine-dependent reductive carboxylation (Fig. S1C).

**Fig. 1.**
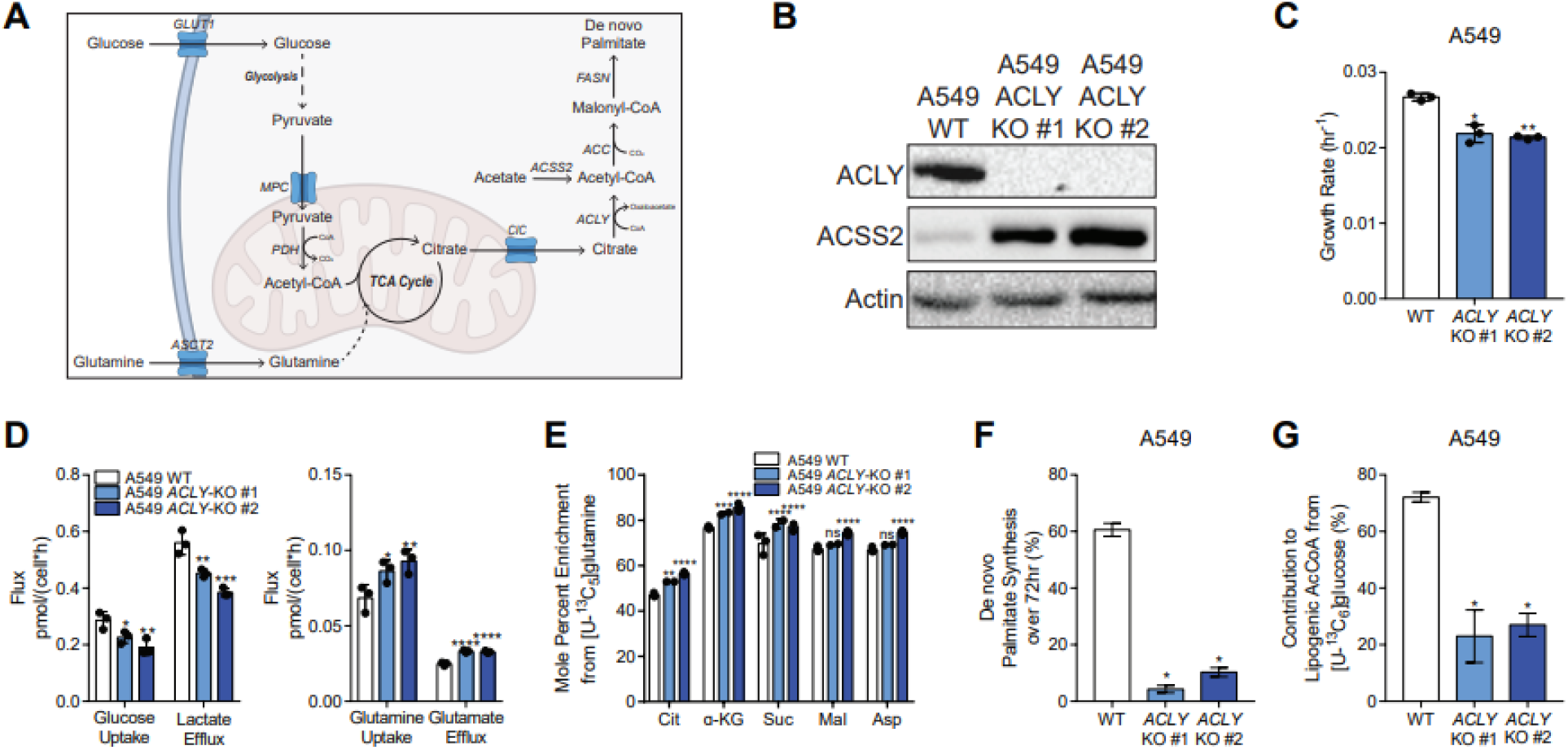
*ACLY*-KO rewires central carbon metabolism and leads to a reduction of palmitate synthesis in cancer cells. (A) The pyruvate-citrate shuttle and fatty acid synthesis. (B) Western Blots of ACLY, ACSS2, and actin in A549 WT and *ACLY*-KO cells. (C) Growth rates of A549 WT and *ACLY*-KO cells grown in high glucose DMEM +10% FBS for 4 days. (D) Glucose uptake, lactate efflux (left) and glutamine uptake, glutamate efflux (right) in A549 WT and *ACLY*-KO cells cultured in high glucose DMEM +10% dFBS for 72 hours (n=3). (E) Mole percent enrichment of TCA intermediates from [U-^13^C_5_]glutamine in A549 *ACLY*-KO clones cultured in high glucose DMEM +10% dFBS for 72 hours (n=3). (F) De novo synthesis of palmitate in A549 *ACLY*-KO cells cultured in [U-^13^C_6_]glucose +10% dFBS for 72 hours (n=3). (G) Percent of lipogenic acetyl-CoA contributed by [U-^13^C_6_]glucose in A549 *ACLY*-KO cells cultured in high glucose DMEM +10% dFBS 72 hours (n=3). In (C-G) data are plotted as mean ± SD. Statistical significance is relative to WT as determined by One-way ANOVA w/ Dunnet’s method for multiple comparisons (C-E) with *, P value < 0.05; **, P value < 0.01; ***, P value < 0.001, ****, P value < 0.0001. In (F,G) data are plotted as mean ± 95% confidence interval (CI). Statistical significance by non-overlapping confidence intervals, *. Unless indicated, all data represent biological triplicates. Data shown are from one of at least two separate experiments. See also Figure S1.

HepG2 cells express the plasma membrane citrate transporter SLC13A5 (*44*), so we next used [2,4-^13^C_2_]citrate to directly trace the ACLY reaction in HepG2 *ACLY*-KO cells,. Catabolism of [2,4-^13^C_2_]citrate provides insight into compartment-specific metabolism as M2 labeling of TCA intermediates will result from direct mitochondrial catabolism of labeled citrate, while M1 isotopologues will be formed if the tracer is first catabolized in the cytosol by ACLY (Fig. S1D). When culturing HepG2 wildtype and *ACLY*-KO cells in the presence of 500 μM [2,4-^13^C_2_]citrate, we found the ratio of M1/M2 labeling on malate, aspartate, succinate, and fumarate was significantly decreased in the *ACLY*-KO cells (Fig. S1E). Furthermore, the relative abundance of M2 alpha-ketoglutarate was comparable to M1 labeling in wildtype cells but was significantly greater in the *ACLY*-KO cells (Fig. S1F). Additionally, enrichment of palmitate was minimal in the *ACLY*-KO cells compared to wildtype (Fig. S1G). These labeling patterns confirm that *ACLY*-KO forces cells to oxidize citrate in the mitochondria and/or cytosol to alpha-ketoglutarate at the expense of acetyl-CoA generation. Next, we applied ISA to quantify palmitate synthesis and the contribution of [U-^13^C_6_]glucose to the lipogenic acetyl-CoA pool. ACLY-deficiency profoundly decreased palmitate synthesis compared to control cells (Fig. 1F, S1H, I). Notably, *ACLY*-KO cells maintained some flux from [U-^13^C_6_]glucose to lipogenic acetyl-CoA such that ~20% of palmitate carbon was derived from glucose in A549 and HepG2 cells, while a smaller decrease was observed in 634T cells (Fig 1G, S1J). Collectively, these results indicate that ACLY deficiency effectively decreases FASN flux and alters cytosolic citrate shuttling under normal culture conditions while reducing growth slightly.

### Acetate metabolism by ACSS2 supports lipogenesis to enable growth of ACLY-deficient cells

Under conditions where glucose-derived citrate is limiting, such as hypoxia, acetate can be “activated” by acetyl-CoA synthetase enzymes. As protein expression of ACSS2 was markedly increased in *ACLY*-KO cells (Fig. 1B, S1A), we hypothesized that acetate utilization may compensate for loss of ACLY. Supplementation of 1 mM [1,2-^13^C_2_]acetate to cultures significantly increased palmitate synthesis in cells lacking ACLY (Fig. 2A, S2A). Acetate became the primary lipogenic substrate in all tested *ACLY*-KO cell lines (Fig. 2B, S2B) but only contributed between 5-40% of lipogenic acetyl-CoA pools in control cells. *De novo* palmitate synthesis was significantly rescued in the *ACLY*-KO cells grown with acetate (Fig. 2A, S2A) compared to those grown in the absence of acetate (Fig. 1F, S1F,G) in all cell lines. We also measured the growth rate of control and ACLY-deficient cell lines in various culture media. *ACLY*-KO cells grew comparably to control cells in media supplemented with 10% dialyzed FBS (dFBS) (Fig 2C, S2C); however, when we removed serum lipids growth of all *ACLY*-KO cells was severely compromised (Fig 2C, S2C). Addition of 1 mM exogenous acetate enhanced the growth of all *ACLY*-KO cell lines while having no impact on growth rates of control cells (Fig 2C, S2C). The most dramatic growth impact of acetate was observed in cultures grown in delipidated serum (Fig 2C, S2C), highlighting the importance of this metabolic redundancy in supporting acetyl-CoA for lipid synthesis (Fig. 2D).

**Fig. 2.**
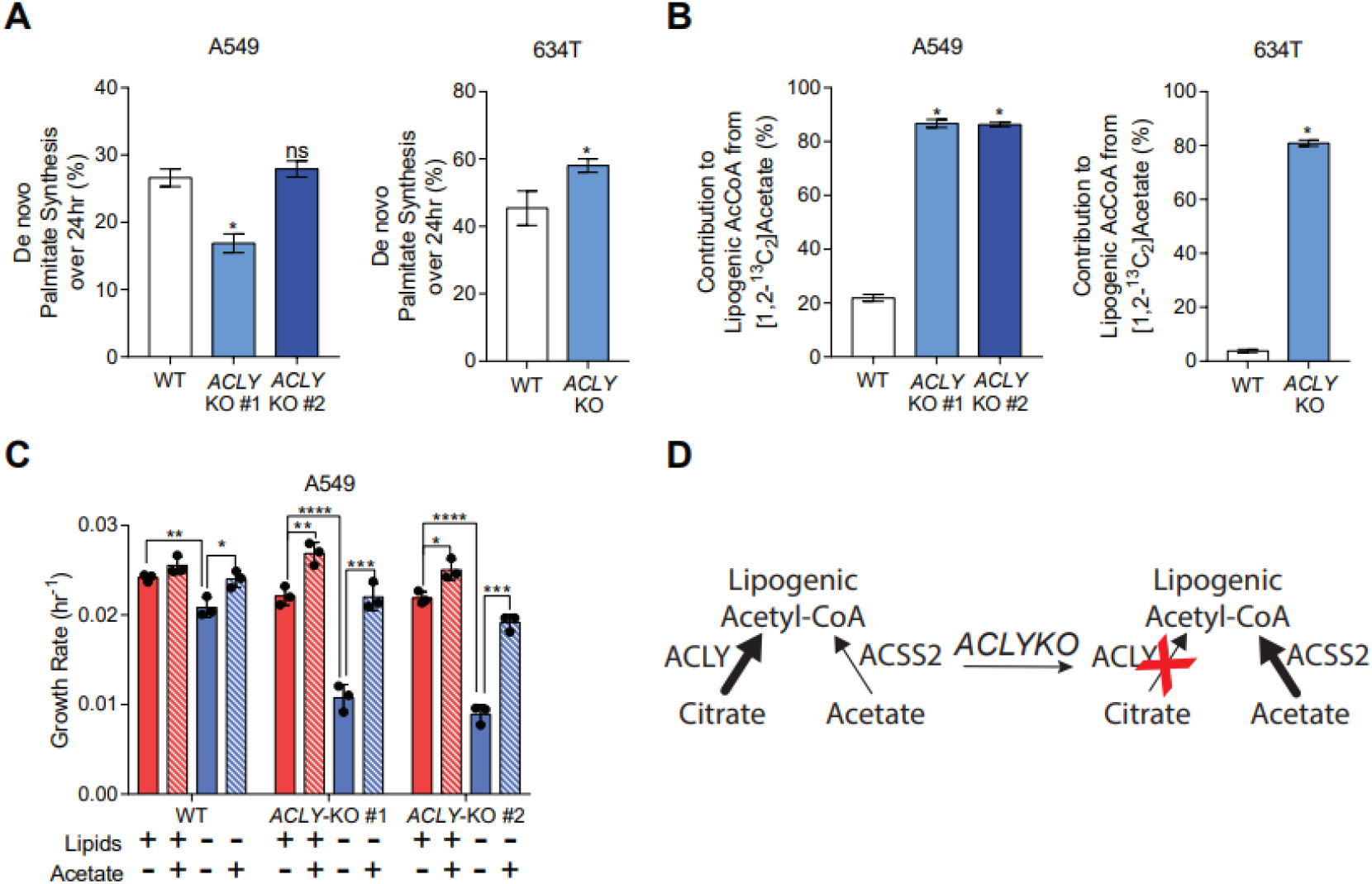
*ACLY*-KO growth and fatty acid synthesis is rescued with addition of extracellular acetate. (A) De novo synthesis of palmitate in A549 (left) and 634T (right) WT and *ACLY*-KO cells cultured in high glucose DMEM +10% dFBS + 1 mM acetate for 24 hours (n=3). (B) Percent of lipogenic acetyl-CoA contributed by [1,2-^13^C_2_]acetate in A549 (left) and 634T (right) and *ACLY*-KO cells cultured in high glucose DMEM +10% dFBS + 1 mM acetate for 24 hours (n=3). (C) Growth rates of A549 WT and *ACLY*-KO cells grown in high glucose DMEM +10% dFBS or delipidated dFBS +/- 1 mM acetate for 4 days (n=3). (D) Schematic of lipogenic acetyl-CoA synthesis in *ACLY*-KO cells. In (A,B) data are plotted as mean ± 95% confidence interval (CI). Statistical significance by non-overlapping confidence intervals, *. In (C) data are plotted as mean ± SD. Statistical significance is determined by Two-way ANOVA w/ Tukey’s method for multiple comparisons with *, P value < 0.05; **, P value < 0.01; ***, P value < 0.001, ****, P value < 0.0001. Unless indicated, all data represent biological triplicates. Data shown are from one of at least two separate experiments. See also Figure S2.

To confirm the role of ACSS2 in supporting lipogenesis, we examined the growth and metabolism of A549 and HepG2 cells lacking ACSS2 (Fig. 3A, S3A). While ACLY expression was unchanged in *ACSS2*-KO cells (Fig. 3A, S3A), the conversion of [1,2-^13^C_2_]acetate to palmitate was significantly decreased (Fig. 3B, S3B), and there was no impact on palmitate synthesis rates in cells lacking ACSS2 (Fig. 3C, S3C). Furthermore, the growth rates of both A549 and HepG2 cell lines were minimally impacted by *ACSS2*-KO in lipid-replete medium (Fig. 3D, S3D). However, HepG2 ACSS2-KO cells observed a significant growth defect compared to wildtype in lipid-depleted media (Fig S3D), matching previous studies which found ACSS2-KD was detrimental to cancer cell growth in conditions of metabolic stress (*31*, *34*). A549 ACSS2-KO cell growth remained unaffected by media delipidation (Fig. 3D). A similar phenotype bifurcation between the cell lines was observed previously, as HepG2 cells exhibited a greater growth rate reduction upon loss of ACLY compared to A549 cells (Fig. 1C, S1B). Interestingly, mRNA expression of ACSS1 and ACSS3, both mitochondrial isozymes of ACSS2, were significantly elevated in HepG2 hepatocarcinoma cells compared to A549 NSCLC cells (Fig. 3E). Furthermore, [1,2-^13^C_2_]acetate enrichment of TCA intermediates was significantly higher in HepG2 cells compared to A549 cells (Fig. 3F,G), suggesting that mitochondrial ACSS1 and/or ACSS3 compete with cytosolic ACSS2 for utilization of acetate and likely reflecting the metabolism in their tissues of origin. This metabolic redundancy may influence acetate shuttling to mitochondria, lipid pathways (e.g. ER), and nucleus to impact epigenetic regulation which is altered by ACSS1/2 knockdown (*45*) but also suggests this pathway may not be targetable in all tumor or cell types.

**Fig. 3.**
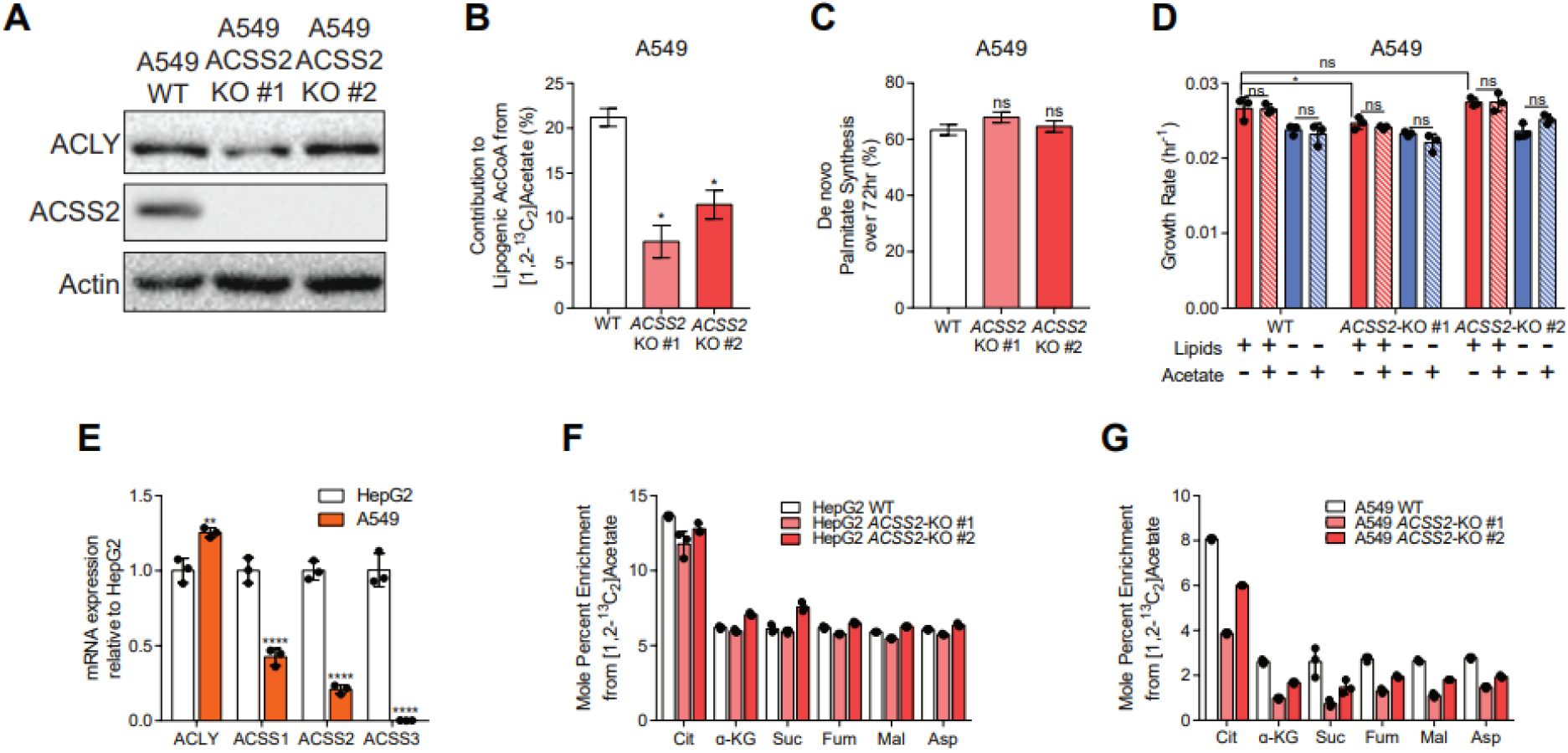
*ACSS2*-KO attenuates catabolism of exogenous acetate with minimal effect on glucose catabolism. (A) Western Blots of ACLY, ACSS2, and actin in A549 WT and *ACSS2*-KO cells. (B) Percent of lipogenic acetyl-CoA contributed by [1,2-^13^C_2_] acetate in A549 *ACSS2*-KO cells cultured in high glucose DMEM +10% dFBS + 1 mM acetate for 24 hours (n=3). (C) De novo synthesis of palmitate in A549 *ACSS2*-KO cells cultured in high glucose DMEM +10% dFBS for 72 hours (n=3). (D) Growth rates of A549 WT and *ACSS2*-KO cells grown in high glucose DMEM +10% dFBS or delipidated dFBS +/- 1 mM acetate for 4 days (n=3). (E) mRNA expression of acetyl-CoA generating enzymes in HepG2 and A549 cells (n=3). (F) Mole percent enrichment of TCA intermediates from [1,2-^13^C_2_] acetate in HepG2 *ACSS2*-KO cells cultured in high glucose DMEM +10% dFBS + 1 mM acetate for 24 hours (n=3). (G) Mole percent enrichment of TCA intermediates from [1,2-^13^C_2_] acetate in A549 *ACSS2*-KO cells cultured in high glucose DMEM +10% dFBS + 1 mM acetate for 24 hours (n=3). In (B, C) data are plotted as mean ± 95% confidence interval (CI). Statistical significance by non-overlapping confidence intervals, *. In (D-G) data are plotted as mean ± SD. Statistical significance is determined by Two-way ANOVA w/ Tukey’s method for multiple comparisons (D); or relative to HepG2 as determined by two-sided Student’s t-test (G) with *, P value < 0.05; **, P value < 0.01; ***, P value < 0.001, ****, P value < 0.0001. Unless indicated, all data represent biological triplicates. Data shown are from one of at least two separate experiments. See also Figure S3.

### Alternative pathways compensate for combined loss of ACLY and ACSS2

Next we characterized A549 ACLY/ACSS2 double knockout (*ACLY*/*ACSS2*-DKO)cells to explore how cancer cells respond to extreme restriction of acetyl-CoA sourcing (Fig. 4A). As expected, growth of A549 *ACLY*/*ACSS2*-DKO cells was significantly impaired relative to wildtype in all conditions tested, and we observed no rescue-effect with acetate supplementation (Fig 4B). We also detected little to no isotope enrichment in palmitate from [U-^13^C_6_]glucose or [1,2-^13^C_2_]acetate in *ACLY*/*ACSS2*-DKO cells (Fig. 4C, D), confirming that both pathways were defective in these cells. Incorporation of polyunsaturated fatty acids (PUFA) into phosphatidylcholine (PC) lipids and triacyl glycerides (TAG) was elevated in *ACLY*-KO and *ACLY*/*ACSS2*-DKO cells, indicating increased reliance on exogenous fatty acids for membrane lipid synthesis in ACLY deficient cells (Fig. 4E, S4A, B). PUFA-containing lipids drive sensitivity to lipid peroxidation (*46*), and we found that PUFA-enriched *ACLY*/*ACSS2*-DKO cells were more susceptible to ferroptosis induced by application of the glutathione peroxidase 4 (GPX4) inhibitor, ML210, than wildtype cells (Fig. 4F). Despite these alterations in lipid metabolism, whole cell protein acetylation was maintained in lipid-replete *ACLY*-KO and *ACLY*/*ACSS2*-DKO cells (Fig 4G, H), suggesting that NSCLC cells are able to sustain acetyl-CoA availability through pathways independent of ACLY and ACSS2.

**Fig. 4.**
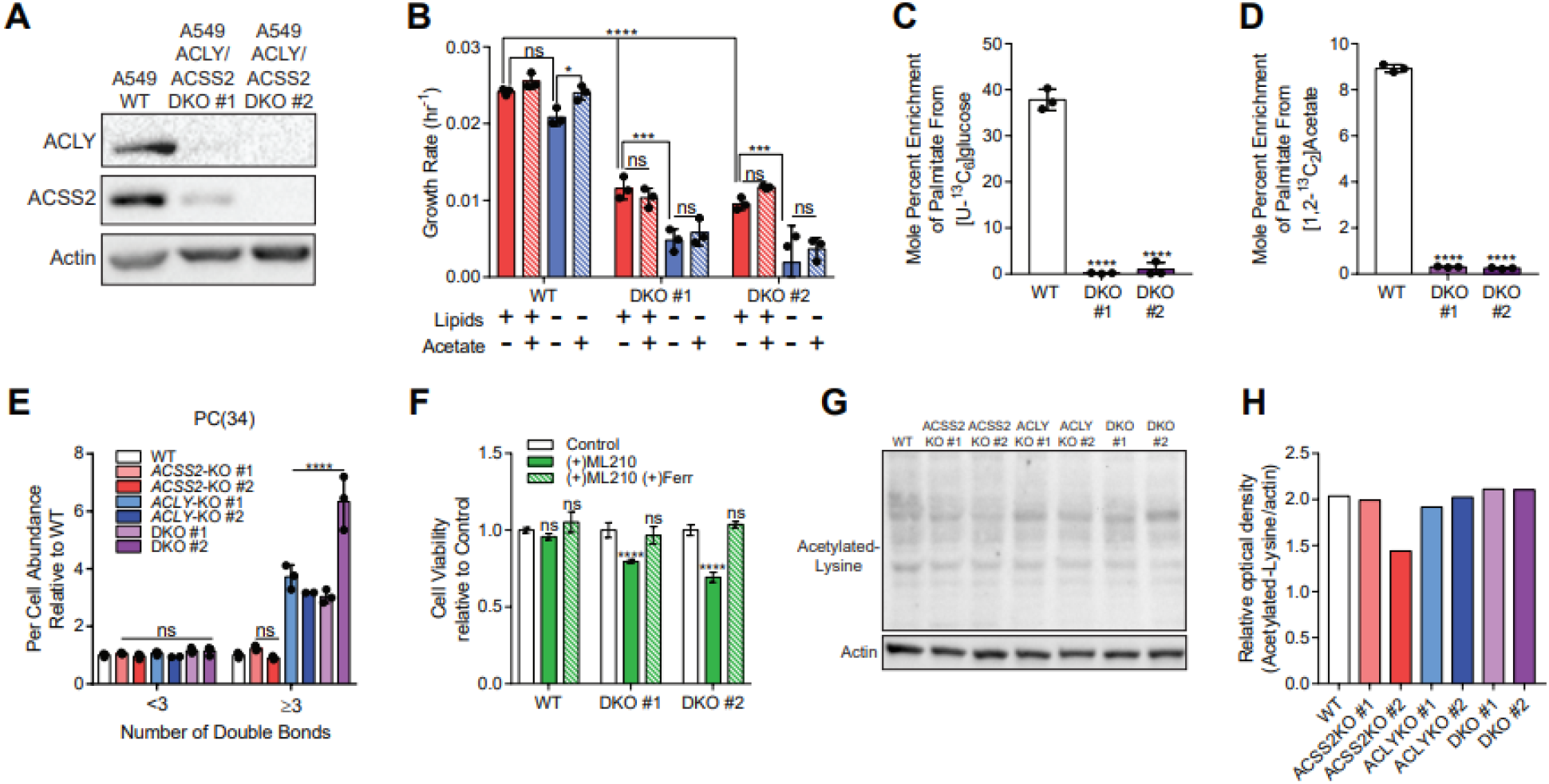
*ACLY*/*ACSS2*-DKO cells are reliant on extracellular lipids, with minimal change in protein acetylation. (A) Western Blots of ACLY, ACSS2, and actin in A549 WT and *ACLY*/*ACSS2*-DKO cells. (B) Growth rates of A549 WT and *ACLY*/*ACSS2*-DKO cells grown in high glucose DMEM +10% dFBS or delipidated dFBS +/- 1 mM acetate for 4 days (n=3). (C) Mole percent enrichment of palmitate from [U-^13^C_6_] glucose in A549 *ACLY*/*ACSS2*-DKO cells cultured in high glucose DMEM +10% dFBS for 72 hours (n=3). (D) Mole percent enrichment of palmitate from [1,2-^13^C_2_] acetate in A549 *ACLY*/*ACSS2*-DKO cells cultured in high glucose DMEM +10% dFBS + 1 mM acetate for 24 hours (n=3). (E) PC(34) abundance of A549 WT, ACSS2-KO, ACLY-KO, ACLY/ACSS2-DKO cells cultured in high glucose DMEM +10% FBS for 2 days (n=3). (F) Cell viability of A549 WT and ACLY/ACSS2-DKO cells cultured in DMEM+10% dFBS +/- 2 μM ML210 and/or 2 μM Ferrostatin for 48 hours (n=4). (G) Western Blots of acetylated-lysine and actin in A549 WT, ACSS2-KO, ACLY-KO, ACLY/ACSS2 DKO whole cell lysates. (H) Relative optical density of acetylated-lysine/actin in A549 WT, ACSS2-KO, ACLY-KO, ACLY/ACSS2-DKO whole cell lysates (n=1). In (B-F) data are plotted as mean ± SD. Statistical significance is determined by Two-way ANOVA w/ Tukey’s method for multiple comparisons (B); or relative to WT as determined by One-way ANOVA w/ Dunnet’s method for multiple comparisons (C-F) with *, P value < 0.05; **, P value < 0.01; ***, P value < 0.001, ****, P value < 0.0001. Unless indicated, all data represent biological triplicates. Data shown are from one of at least two separate experiments. See also Figure S4.

The continued availability of acetyl-CoA for histone acetylation suggests cancer cells can employ alternate pathways for acetyl-CoA generation under conditions of metabolic stress, including non-enzymatic mechanisms (*47*). To explore these pathways more deeply we cultured A549 *ACLY*-KO and *ACLY*/*ACSS2*-DKO cells with either ^13^C-labeled glucose, glutamine, and pyruvate for 72 hours and quantified isotope enrichment in palmitate. In theory, knockout of ACLY will eliminate glucose to palmitate carbon transfer (Fig. 5A); however, A549 *ACLY*-KO cells traced with [U-^13^C_6_]glucose continued to sustain labeling on newly synthesized palmitate (Fig. 5B), and we observed similar results with [3-^13^C]pyruvate (Fig. 5C). Culture of the *ACLY*-KO and *ACLY*/*ACSS2*-DKO cells with [U-^13^C_5_]glutamine yielded insignificant enrichment of palmitate (Fig. 5D), suggesting the existence of a citrate/ACLY-independent lipogenic pathway which allows glucose and pyruvate but not glutamine to generate acetyl-CoA. We previously demonstrated that carnitine acetyl-transferase (CRAT) actively exports mitochondrial branched-chain-CoAs to enable their elongation to long-chain monomethyl branched-chain fatty acids in adipocytes (*48*), and Izzo et al. have observed that CRAT facilitates nuclear acetylation and lipogenesis (*49*). However, we hypothesize that this activity is insufficient to support lipid synthesis since we observed minimal labeling of palmitate from [U-^13^C_6_]glucose in the A549 *ACLY*/*ACSS2*-DKO (Fig. 5A), despite noting significant enrichment of pyruvate and citrate (Fig. S5A,B). Likewise, ^13^C pyruvate, and glutamine also yielded negligible label on palmitate in A549 *ACLY*/*ACSS2*-DKO cells (Fig. 5B, C, D), suggesting that ACSS2 is involved in the conversion of pyruvate to acetyl-CoA (Fig. 5A). Such a mechanism has been described whereby reactive oxygen species facilitate oxidative decarboxylation of pyruvate to acetate (*50*). The combination of KO cells and tracers applied here confirm this activity supports acetyl-CoA generation, but the extracellular acetate, ACSS2, and lipid dependency of *ACLY*-KO cell growth suggests this *de novo* acetate flux is not significant enough to sustain the biosynthetic needs of the ACLY deficient cells tested here (Fig. 2C, S2C).

**Fig. 5.**
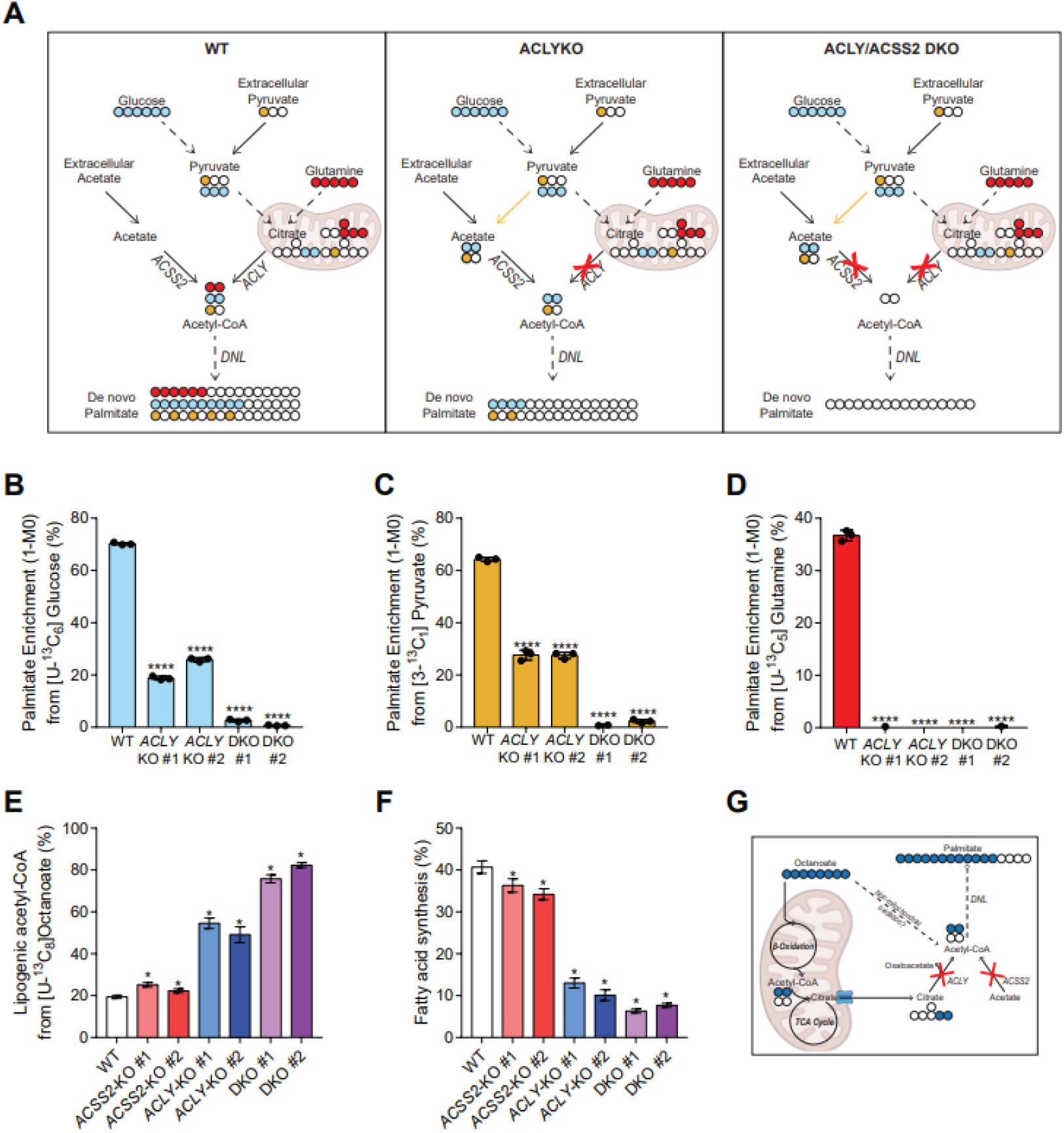
Disruption of canonical acetyl-CoA synthesis induces alternative synthesis pathways. (A) Schematic showing de novo acetate synthesis in *ACLY*-KO cells. Dashed lines represent multiple reactions. (B) Enrichment (1-M0) of palmitate from [U-^13^C_6_] glucose in A549 WT, *ACLY*-KO, *ACLY*/*ACSS2*-DKO cells cultured in high glucose DMEM +10% dFBS for 72 hours (n=3). (C) Enrichment (1-M0) of palmitate from [3-^13^C_1_] pyruvate in A549 WT, *ACLY*-KO, *ACLY*/*ACSS2*-DKO cells cultured in high glucose DMEM +10% dFBS + 5mM pyruvate for 72 hours (n=3). (D) Enrichment (1-M0) of palmitate from [U-^13^C_5_] glutamine in A549 WT, *ACLY*-KO, *ACLY*/*ACSS2*-DKO cells cultured in high glucose DMEM +10% dFBS for 72 hours (n=3). (E) Percent of lipogenic acetyl-CoA contributed by [U-^13^C_8_] octanoate in A549 WT, ACSS2-KO, ACLY-KO, ACLY/ACSS2-DKO cells cultured in high glucose DMEM +10% dFBS + 500 μM octanoate for 24 hours (n=3). (F) De novo synthesis of palmitate in A549 WT, ACSS2-KO, ACLY-KO, ACLY/ACSS2-DKO cells cultured in high glucose DMEM +10% dFBS + 500 μM octanoate for 24 hours (n=3). (G) Schematic of non-mitochondrial acetyl-CoA synthesis from [U-^13^C_8_] octanoate in ACLY/ACSS2-DKO cells. Filled circles are 13C, empty circles are 12C. In (B-D) data are plotted as mean ± SD. Statistical significance relative to WT as determined by One-way ANOVA w/ Dunnet’s method for multiple comparisons (B-F) with *, P value < 0.05; **, P value < 0.01; ***, P value < 0.001, ****, P value < 0.0001. In (F,G) data are plotted as mean ± 95% confidence interval (CI). Statistical significance by non-overlapping confidence intervals, *. Unless indicated, all data represent biological triplicates. Data shown are from one of at least two separate experiments. See also Figure S5.

To examine how oxidation of fatty acyl-CoA might contribute to acetyl-CoA homeostasis we traced the A549 cell panel with 500 μM [U-^13^C_8_]octanoate, which passively diffuses across the mitochondrial membrane as well as peroxisomal membranes (*39*). Notably, octanoate contributed significantly to fatty acid biosynthesis, even in *ACLY*/*ACSS2*-DKO cells (Fig. 5E,F, S5C) and contrasting results with glucose, acetate, or glutamine tracers (Fig 4C, D, 5D). *ACLY*-KO and *ACLY*/*ACSS2*-DKO cells had a 2.5-fold and 4-fold increase, respectively, in enrichment of the lipogenic acetyl-CoA pool from octanoate compared to the wildtype or *ACSS2*-KO cells (Fig. 5E). Similar trends in lipogenic acetyl-CoA labeling from [U-^13^C_8_]octanoate were observed in HepG2 and 634T *ACLY*-KO clones (Fig. S5C). Indeed, supplementation of octanoate enabled quantitation of fractional new fatty acid synthesis in *ACLY*/*ACSS2*-DKO cells by ISA (Fig. 5F). Citrate enrichment from octanoate was also increased in the *ACLY*-KO and *ACLY*/*ACSS2*-DKO cells (Fig. S5D). However, based on the previous [U-^13^C_5_]glutamine tracing results (Fig. 5D) mitochondrial citrate minimally contributes to fatty acid synthesis in the *ACLY*-KO or *ACLY*/*ACSS2*-DKO cells, and no carnitine was added to cultures(*49*), suggesting that ACLY-deficient cells use peroxisomal metabolism to supply acetyl-CoA for biosynthesis (Fig 5G).

### Peroxisomal β-oxidation supplies lipogenic acetyl-CoA in the absence of ACLY and ACSS2

Peroxisomes are the site of β-oxidation for very long chain fatty acids (VLCFAs) and branched chain fatty acids (BCFAs). Peroxisomes are closely associated with lipid droplets and contain the metabolic machinery for acetyl-CoA generation from diverse fatty acids (*51*). In fact, Izzo et al. observed increased expression of peroxisomal fatty acid oxidation genes in their *ACLY*/*ACSS2*-DKO cell lines (*49*). To test whether peroxisomes could serve as an appreciable source of acetyl-CoA in cancer cells, we profiled lipid abundances in the A549 cell panel. While total TAGs were slightly reduced in most cells, total PCs were unchanged (Fig. 6A). However, we detected significant alterations in the acyl-chain composition of the lipids, such that TAGs and PCs containing VLCFAs were depleted in *ACLY*-KO and *ACLY*/*ACSS2*-DKO cells (Fig. 6B, C). This depletion is complete for nearly all measurable VLCFA containing saturated, mono-, and di-unsaturated TAG species (Fig. S6A) as well as saturated, and mono-unsaturated PC species (Fig S6B). Depletion of VLCFAs in lipids suggests altered VLCFA homeostasis, potentially due to increased VLCFA oxidation in the peroxisome or decreased elongation using acetyl-CoA.

**Fig. 6.**
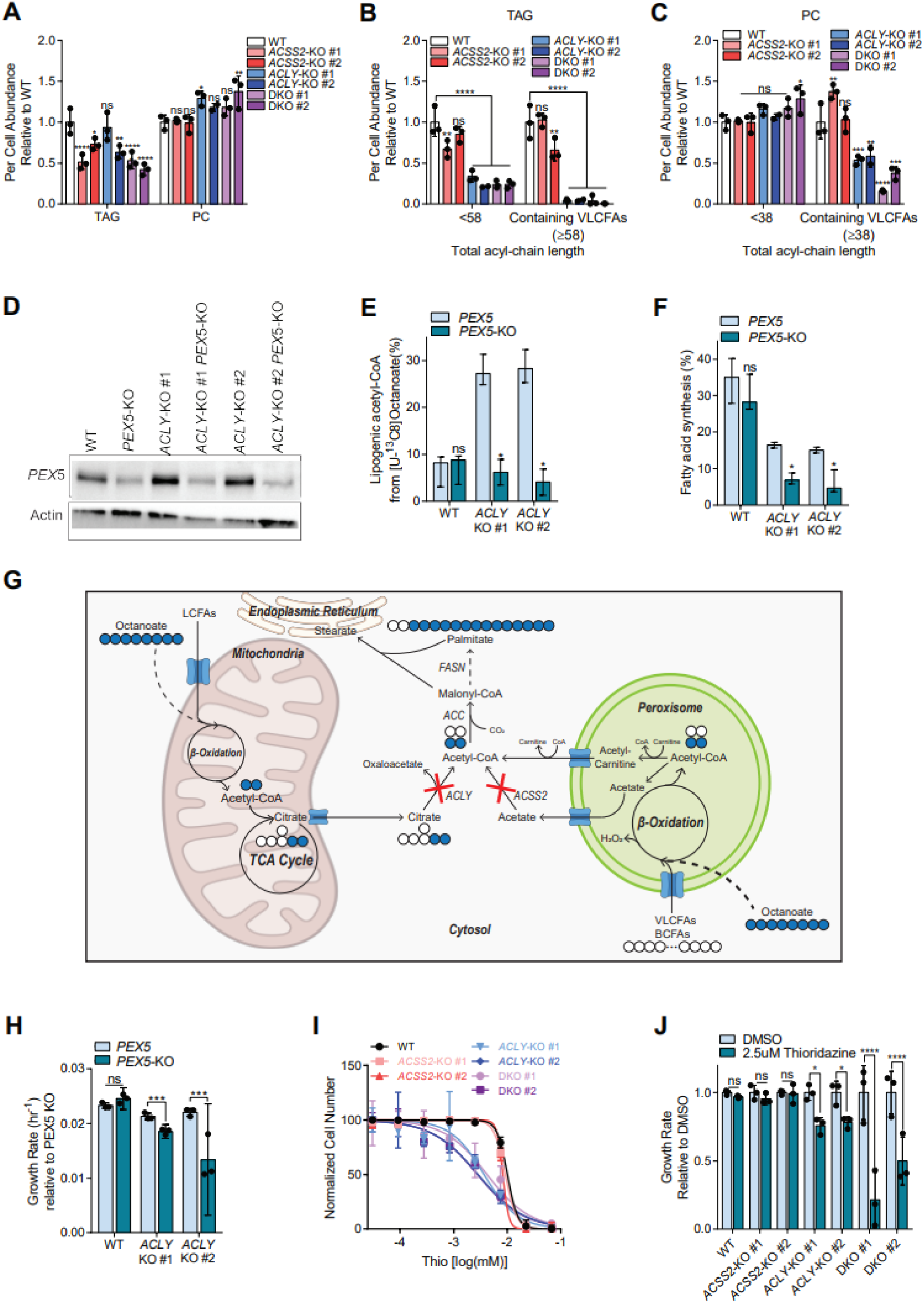
Peroxisomal β-oxidation becomes a major source of lipogenic acetyl-CoA with ACLYKO and ACLY/ACSS2 DKO. (A) Total abundance of TAGs and PCs in A549 WT, ACSS2-KO, ACLY-KO, ACLY/ACSS2-DKO cells cultured in high glucose DMEM +10% FBS for 2 days (n=3). (B) Total abundance of saturated, mono- and di-unsaturated TAGs per cell of A549 WT, ACSS2-KO, ACLY-KO, ACLY/ACSS2-DKO cells cultured in high glucose DMEM +10% FBS for 2 days (n=3). (C) Total abundance of saturated and monounsaturated PCs per cell of A549 WT, ACSS2-KO, ACLY-KO, ACLY/ACSS2-DKO cells cultured in high glucose DMEM +10% FBS for 2 days (n=3). (D) Western blots of PEX5 and actin in A549 WT, PEX5 KO, ACLY KO and ACLY-PEX5-KO cells. (E) Percent of lipogenic acetyl-CoA contributed by [U-^13^C_8_]] octanoate in A549 WT, PEX5-KO, ACLY-KO and ACLY/PEX-5 KO cells cultured in high glucose DMEM +10% dFBS for 48 hours (n=3). (F) De novo synthesis of palmitate in A549 WT, PEX5-KO, ACLY-KO and ACLY/PEX-5 KO cells cultured in high glucose DMEM +10% dFBS for 72 hours (n=3). (G) Schematic of peroxisomal acetyl-CoA synthesis from [U-^13^C_8_] octanoate in *ACLY*/*ACSS2*-DKO cells. Filled circles are 13C, empty circles are 12C. (H) Growth rates of A549 WT and ACLY-KO relative to PEX5 KO cells grown in high glucose DMEM +10% FBS for 4 days. (I) Dose response of A549 WT, *ACSS2*-KO, *ACLY*-KO, *ACLY*/*ACSS2*-DKO cells cultured in high glucose DMEM +10% dFBS to Thioridazine over 4 days (n=3). (J) Growth rates of A549 WT and ACLY/ACSS2-DKO cells grown in high glucose DMEM +10% dFBS +/- 2.5 μM thioridazine relative to DMSO for 4 days (n=3). In (A-C, H, J) data are plotted as mean ± SD. Statistical significance as determined by One-way ANOVA w/ Dunnet’s method for multiple comparisons relative to WT (A-C) or DMSO (J) with *, P value < 0.05; **, P value < 0.01; ***, P value < 0.001, ****, P value < 0.0001. In (E,F) data are plotted as mean ± 95% confidence interval (CI). Statistical significance by non-overlapping confidence intervals, *. Unless indicated, all data represent biological triplicates. Data shown are from one of at least two separate experiments. See also Figure S6.

To further investigate we studied role of peroxisomes in non-canonical lipogenic acetyl-CoA contribution. Peroxisome maintenance depends on the import of nuclear-encoded proteins containing a peroxisomal targeting signals (PTS1), which are recognized by cytosolic PEX5 receptors that transport them into the peroxisomal lumen. In the absence of PEX5 peroxisomal enzyme transport is ablated and metabolism altered (*52*). To further investigate the role of peroxisomes in supporting acetyl-CoA metabolism we generated *PEX5* and *ACLY*/*PEX5*-knockout (*ACLY*/*PEX5*-KO) clones using CRISPR/Cas9 in A549 cells (Fig. 6D). *ACLY*/*ACSS2*/*PEX5* triple knockout cells did not proliferate and could not be tested. We then cultured A549 *PEX5*-KO cells with 500 μM [U-^13^C_8_]octanoate and quantified its contribution to the lipogenic acetyl-CoA pool. Intriguingly, both enrichment from octanoate and fatty acid synthesis were significantly lower in *ACLY*/*PEX5*-KO cells compared to *ACLY*-KO cells (Fig 6E-F), providing evidence that peroxisomal metabolism supports acetyl-CoA and lipid synthesis. On the other hand, we saw no effect of PEX5 deficiency on lipogenic acetyl-CoA enrichment or palmitate synthesis in wildtype cells (Fig 6E-F). The labeling distribution of palmitate from [U-^13^C_8_]octanoate highlights these changes, as *ACLY*-KO cells showed robust incorporation but *ACLY*/*PEX5*-KO cells showing predominantly M8 palmitate (or direct incorporation of octanoate; Fig. S6C). These data suggest that knockout of PEX5 significantly disrupts the contribution of peroxisomal fatty acid oxidation to lipogenic acetyl-CoA production in the ACLY-KO cells (Fig 6G). In fact, we also observed a decrease in octanoate-mediated histone acetylation in *ACLY*/*PEX5*-KOs compared to *ACLY*-KO cells in delipidated media (Fig. S6D).

To elucidate how impairing peroxisomal β-oxidation affects proliferation of cells lacking ACLY, we quantified the growth of each cell line in response to knockout of *PEX5*. Strikingly, *ACLY*/*PEX5*-KO cells showed a significant decrease in growth relative to *ACLY*-KO cells, whereas there was no growth effect on single *PEX5*-KO cells compared to wildtype (Fig. 6H). Finally, we quantified the growth the entire A549 ACLY/ACSS2-KO cell panel in response to increasing concentrations of the peroxisomal β-oxidation inhibitor, Thioridazine (Thio) (*53*, *54*) (Fig 6I). Strikingly, *ACLY*-KO and *ACLY*/*ACSS2*-DKO cells were significantly more sensitive to Thio treatment than wildtype or *ACSS2*-KO cells (Fig. 6J). Application of 2.5 μM Thio had no appreciable impact on growth of wildtype and *ACSS2*-KO cells, but significantly decreased proliferation of *ACLY*-KO and *ACLY*/*ACSS2*-DKO cells (Fig. 6I). Together, these results suggest that peroxisomal β-oxidation provides a significant growth sustaining level of acetyl-CoA in the absence of functional ACLY. Multiple mechanisms have been suggested for shuttling peroxisomal acetyl-CoA to the cytosol in mammalian cells, including conversion to acetate by a thioesterase and subsequent export (and activation by ACSS2 in the cytosol) as well as shuttling as acetyl-carnitine facilitated by peroxisomal CRAT (*55*–*57*). The elevated palmitate enrichment from [U-^13^C_8_]octanoate observed in *ACLY*/*ACSS2*-DKO cells suggests that these cells employ an acetate-independent acetyl-CoA export mechanism, potentially via activity of peroxisomal CRAT (Fig. 6G), which is also localized to mitochondria.

## Discussion

While the core metabolic machinery of cancer cells acts “as advertised” under nutrient replete conditions, the diverse stresses experienced the tumor microenvironment induce complex rewiring of metabolic fluxes between organelles, including mitochondria, the ER, peroxisomes, and the plasma membrane. Here we show this organelle crosstalk is critical for cancer cells to maintain acetyl-CoA homeostasis and metabolic resilience when organelle-specific pathways become compromised. Acetyl-CoA serves as a substrate regulating numerous processes across mitochondria, the nucleus, peroxisomes, the ER, and plasma membrane but importantly cannot cross membranes. As such, understanding how acetyl-groups are shuttled throughout the cell for bioenergetics, biosynthesis, and cellular regulation is critically important. Here we applied metabolic tracing and lipogenesis studies to cancer cell lines where key pathways for acetyl-CoA generation were knocked out, including ACLY, ACSS2, and peroxisomal metabolism (PEX5). We further highlight a key role for peroxisomal metabolism in providing acetyl-CoA for fatty acid synthesis and histone acetylation.

Acetyl-CoA-fueled fatty acid synthesis is important for cancer progression in NSCLC models, whether this occurs in endogenous tissues or in situ within tumors (*7*). Targeting of ACLY to reduce tumor cell growth and increase dependence on exogenous lipids has also been explored in detail (*24*–*26*, *58*). Likewise, ACSS2 has garnered significant interest as a mechanism through which cancer cells source acetyl-CoA for growth and survival (*31*–*33*, *59*). ACSS2 expression and acetate-dependent flux both increase in the context of ACLY deficiency (*36*), and trafficking of acetate to the nucleus for histone acetylation may be an important aspect of this enzyme’s function (*45*, *60*–*62*). The impact of ACSS isozyme expression on acetate shuttling, fatty acid synthesis and epigenetic regulation warrants further study as it may provide insight into how therapeutic interventions targeting acetate catabolism impact whole body physiology. Peroxisomes further complicate these questions, particularly in reference to their impact on lipid homeostasis and diversity. Understanding how these nodes of metabolic exchange are exploited by cancer cells shed light on resistance mechanisms, as these redundancies in pathway architecture can reduce efficacy of drugs targeting metabolic enzymes (*63*).

Beyond cancer, understanding how cells manage intracellular metabolic pools using these KO and tracing approaches is also relevant for states of “overnutrition.” Indeed, exogenous treatment of fatty acids or lipids to cells in vitro often occurs at supra-physiological levels, so our studies further highlight the role of peroxisomal metabolism in generating acetyl-CoA for lipid biosynthesis and acetylation (*51*). While these KO systems lack some physiology in the acute nature of pathway targeting, they also allow one to focus biochemical studies on these alternate pathways when substrate levels are high. However, we and Luke Izzo et al(*49*). in this issue also note key questions that remain such as the relative contribution of mitochondrial versus peroxisomal oxidation and the dependence on carnitine for transport since both organelles express CRAT (*64*, *65*). The relative contribution of each pathway likely shifts as a function of cell and tissue type, and these pathways may also be distinct with respect to how they contribute to downstream lipid synthesis versus histone acetylation. By understanding how these metabolic pathways are balanced they can be more effectively targeted both in neoplastic or metabolically stressed tissues.

## Materials and Methods

### Cell Lines

A549 and HepG2 cells were obtained from HDBiosciences, HAP1 cells were obtained from Horizon Discovery, and 634T cells were obtained from the Shaw Lab at the Salk Institute for Biological Sciences. All cell lines were incubated at 37C with 5% CO_2_ and cultured using Dulbecco’s Modified Eagle Media (DMEM) with 10% Fetal Bovine Serum and 1% Penicillin-Streptomycin. Cells tested negative for mycoplasma contamination. All media were adjusted to pH = 7.3.

### Cell Proliferation and ^13^C Tracing

Proliferation studies were performed on 12 well plates with an initial cell number of 50,000/well for A549s and 100,000/well for HepG2s. Cells were plated in growth media and allowed to adhere for 24 hours before changing to the specified growth media. Cell counts were performed at days 0 and 4 using a hemocytometer.

^13^C isotope tracing media was formulated using glucose and glutamine free DMEM 5030 supplemented with either 20 mM [U-^13^C_6_]glucose, 4 mM [U-^13^C_5_]glutamine, 1 mM [1,2-^13^C_2_]acetate, 5 mM [3-^13^C_1_]pyruvate, 500 μM [U-^13^C_8_] octanoate or 500 μM [2,4-^13^C_2_]citrate (Cambridge Isotopes) and 10% dialyzed FBS. ^13^C palmitate studies were performed with 200 μM albumin conjugated [U-^13^C_16_] palmitate and 10% dialyzed delipidated FBS. All studies were performed with a final concentration of 20 mM glucose and 4 mM glutamine. Cultured cells were washed with 1 mL PBS prior to applying tracing media for 6-72 hours as indicated in figure legends.

### Drug Dose Response

Drug dose response studies were performed on 96 well plates with an initial cell number of 2,500/well. Cells were plated in growth media and allowed to adhere for 24 hours before changing to the specified growth media with the indicated concentration of thioridazine. After 4 days wells were washed 2x with tap water, 50uL of 0.5% crystal violet staining solution were added to each well for 20 minutes. Crystal violet stain was removed and wells were washed 3x with tap water and air dried overnight. Next, 200 uL methanol was added to each well and plates were incubated at room temperature for 20 minutes. Optical density of wells were measured at 570 nm on a TECAN plate reader.

### Ferroptosis Assay

Wildtype and ACLY/ACSS2-DKO cells were plated in 96 well plates with an initial cell number of 5,000/well. Cells were plated in growth media and allowed to adhere for 24 hours before changing to the specified growth media with 2uM ML210 and/or 6uM Ferrostatin. After 48 hours, cell viability was quantified using the PrestoBlue™ Cell Viability (Thermo, A13261) and a TECAN plate reader.

### Isotopomer Spectral Analysis (ISA)

Isotopomer spectral analysis (ISA) was performed to estimate the percent of newly synthesized palmitate as well as the contribution of a tracer of interest to the lipogenic acetyl-CoA pool (*66*, *67*). Parameters for contribution of ^13^C tracers to lipogenic acetyl-CoA (D value) and percentage of newly synthesized fatty acid (g(t) value) and their 95% confidence intervals are then calculated using best-fit model from INCA MFA software. Experimental palmitate labeling from [U-^13^C_6_]glucose, [U-^13^C_5_]glutamine, [1,2-^13^C_2_]acetate, [U-^13^C_8_] octanoate or [2,4-^13^C_2_]citrate after a 24-72 hour trace, as indicated in figure legends, was compared to simulated labeling using a reaction network where C16:0 is condensation of 8 AcCoA. ISA data plotted as mean ± 95% CI. * indicates statistical significance by non-overlapping confidence intervals.

### CRISPR/cas9 engineered knockout cell lines

HDBiosciences generated A549 and HepG2 *ACLY*-KO, *ACSS2*-KO, and *ACLY*/*ACSS2*-DKO cell lines. *ACLY*, *ACSS2*, and *ACLY*/*ACSS2* knockout clones were generated using the strategy described previously (*68*). Briefly, a tandem guide RNAs (gRNAs) was designed to target the human *ACLY* (gRNA sequences: gaccagctgatcaaacgtcg, ggggtcaggatgaacgtgtg) and *ACSS2* (gRNA sequences: ctcgcggtagcgctgcagcg, aatggaaaagggattccggg) using the online CRISPR guide tool provided by the Zhang Lab at MIT (http://tools.genome-engineering.org). The gRNA duplex was cloned into lentiCRISPRv2 (Addgene #52961) (*69*). A549 and HepG2 cells were transfected with the *ACLY* and/or *ACSS2* specific gRNA to generate pooled knockouts. After puromycin selection, single-cell clones were isolated by diluting the pooled knockout lines at 1 cell/100 μL and plating 100 μL into each well of a 96 well plate. Clones were maintained by exchanging media every 3-5 days. Clones were validated by PCR and western blot. PEX5 and ACLY/PEX5 knockout clones were generated using the strategy described previously (*70*). Briefly, chemically modified guide RNAs (gRNAs) (Synthego) were designed to target the human PEX5 (gRNA sequences: guugguggcugaauucc). gRNAs were complexed with recombinant Cas9 (IDT Cat#1081059) and introduced into cells by electroporation (4D-Nucleofector, Lonza Biosciences) according to manufacturer protocols. Single cell clones were generated by serial dilution of the bulk knockout population in 96 well plates. Clones were maintained by exchanging the media every 3 days and validated by western blot.

### Lentivirus Production

One 10cm dish of HEK293FT cells at 60% confluency were transfected with 1.3 μg VSV.G/pMD2.G, 5.3 μg lenti-gag/pol/pCMVR8.2, and 4 μg of the gRNA duplexed lentiCRISPRv2 using 16 μL Lipofectamine 3000 diluted in 0.66 mL of OPTI-MEM. Medium containing viral particles was harvested 48 and 72 hours after transfection, then concentrated by Centricon Plus-20 100,000 NMWL centrifugal ultrafilters, divided into aliquots and frozen at −80°C.

### Metabolic Flux Analysis

Metabolic fluxes for citrate were calculated by collecting media at time 0 and spent media after 72 hours. Spent media was centrifuged at 300g for 5 min, to remove cell debris. Cell counts were performed at time 0 and after 72 hours as well.

### Metabolite Extraction and GC-MS Analysis

At the conclusion of the tracer experiment, media was aspirated. Then, cells were rinsed twice with 0.9% saline solution and lysed with 250 μL ice-cold methanol. After 1 minute, 100 μL water containing 1 μg/ml norvaline was added to each sample and vortexed for one minute. 250 μL chloroform was added to each sample, and all were vortexed again for 1 minute. After centrifugation at 21,130 g for 10 minutes at 4°C, 250 μL of the upper aqueous layer was collected and evaporated under vacuum at 4°C. Then, 250 μL of the lower organic layer was collected and evaporated under air at room temperature. Dried polar and nonpolar metabolites were processed for gas chromatography (GC) mass spectrometry (MS) as described previously in Cordes and Metallo (*67*). Briefly, polar metabolites were derivatized using a Gerstel MultiPurpose Sampler (MPS 2XL). Methoxime-tBDMS derivatives were formed by addition of 15 μL 2% (w/v) methoxylamine hydrochloride (MP Biomedicals, Solon, OH) in pyridine and incubated at 45°C for 60 minutes. Samples were then silylated by addition of 15 μL of N-tert-butyldimethylsily-N-methyltrifluoroacetamide (MTBSTFA) with 1% tert-butyldimethylchlorosilane (tBDMS) (Regis Technologies, Morton Grove, IL) and incubated at 45°C for 30 minutes. Nonpolar metabolites were saponified and transesterified to fatty acid methyl esters (FAMEs) by adding 500 μL of 2% H2SO4 in methanol to the dried nonpolar layer and heating at 50°C for 1 hour. FAMEs were then extracted by adding 100 μL of a saturated salt solution and 500 μL hexane and vortexing for 1 minute. The hexane layer was removed, evaporated, and resuspended with 60μL hexane for injection.

Derivatized polar samples were injected into a GC-MS using a DB-35MS column (30m x 0.25mm i.d. x 0.25μm, Agilent J&W Scientific, Santa Clara, CA) installed in an Agilent 7890B GC system integrated with an Agilent 5977a MS. Samples were injected at a GC oven temperature of 100°C which was held for 1 minute before ramping to 255°C at 3.5°C/min then to 320°C at 15°C/min and held for 3 minutes. Electron impact ionization was performed with the MS scanning over the range of 100-650 m/z for polar metabolites.

Derivatized nonpolar samples were injected into a GC-MS using a Fame Select column (100m x 0.25mm i.d. x 0.25μm, Agilent J&W Scientific, Santa Clara, CA) installed in an Agilent 7890A GC system integrated with an Agilent 5977A MS. Samples were injected at a GC oven temperature of 80°C which was held for 1 minute before ramping to 170°C at 20°C/min then to 188°C at 1°C/min then to 250°C at 20°C/min and held for 10 minutes. Electron impact ionization was performed with the MS scanning over the range of 54-400 m/z for nonpolar metabolites.

Metabolite levels and mass isotopomer distributions were analyzed with an in-house Matlab script which integrated the metabolite fragment ions and corrected for natural isotope abundances. Mole percent enrichment calculations were performed using Escher-Trace (*71*).

### LC-MS/MS analysis

Lipids were extracted from confluent 6 well plates after growth in DMEM + 10% FBS + 1% penicillin streptomycin for 48 hours. At the conclusion of the experiment, media was aspirated, and the cells were rinsed twice with saline solution. Cells were lysed with 750 μL of ice cold 1:1 methanol/water solution for 5 minutes on ice and scraped into Eppendorf tubes. Next, 500 μL of ice cold chloroform and 50 uL butylated hydroxytoluene (1mg/mL in methanol) were added to the tube, along with EquiSLASH (Avanti, Croda International Plc, 330731internal standard. The tubes were vortexed for 5 min and centrifuged at 21,230g at 4°C for 5 min. The lower organic phase was collected and 2 μL of formic acid was added to the remaining polar phase which was re-extracted with 500uL of chloroform. Combined organic phases were dried under nitrogen and the pellet was resuspended in 50 μL isopropyl alcohol.

Q Exactive orbitrap mass spectrometer with a Vanquish Flex Binary UHPLC system (Thermo Scientific) was used with an Accucore C30, 150 x 2.1mm, 2.6 μm particle (Thermo) column at 40°C. 5 μL of sample was injected. Chromatography was performed using a gradient of 40:60 v/v water: acetonitrile with 10 mM ammonium formate and 0.1% formic acid (mobile phase A) and 10:90 v/v acetonitrile: 2-propanol with 10 mM ammonium formate and 0.1% formic acid (mobile phase B), both at a flow rate of 0.2 mL/min. The LC gradient ran from 30% to 43% B from 3-8 min, then from 43% to 50% B from 8-9min, then 50-90% B from 9-18min, then 90-99% B from 18-26 min, then held at 99% B from 26-30min, before returning to 30% B in 6 min and held for a further 4 min.

Lipids were analyzed in positive mode using spray voltage 3.2 kV. Sweep gas flow was 1 arbitrary units, auxiliary gas flow 2 arbitrary units and sheath gas flow 40 arbitrary units, with a capillary temperature of 325°C. Full MS (scan range 200-2000 m/z) was used at 70 000 resolution with 1e6 automatic gain control and a maximum injection time of 100 ms. Data dependent MS2 (Top 6) mode at 17 500 resolution with automatic gain control set at 1e5 with a maximum injection time of 50 ms was used. Data was analyzed using EI-Maven and Escher-Trace software, and peaks normalized to Avanti EquiSPLASH internal standard. Lipid species specific fragments used for identification and quantitation are presented in the Supplementary Table 1

### Western Blot

A549, HepG2, HAP1, and 634T wildtype, ACLY and/or ACSS2-KO cell lines were lysed in M-PER buffer (Thermo Scientific, 78501) with 1x protease inhibitor (Sigma-Aldrich). Protein concentrations were determined using Pierce^™^ BCA protein assay kit (Thermo Scientific, 23225). 15 ug protein were loaded and separated using 4–15% Mini-PROTEAN^®^ TGX^™^ Precast SDS-PAGE Gels (Bio-rad, #4561086). Samples were then transferred to nitrocellulose membranes for immunoblotting. Total acetylated-lysine (Cell Signaling Technology, 9441), β-Actin (Sigma, A5441), ACLY (Cell Signaling Technology, 13390), and ACSS2 (Cell Signaling Technology, 3658), PEX5(D7V4D) (Cell Signaling Technology, 83020) antibodies were used to probe their respective targets. Anti-Rabbit horseradish peroxidase-conjugated secondary antibodies (Millipore, AP132P) were used for imaging.

### RNA isolation and quantitative RT-PCR

Total RNA was purified from cultured cells using Trizol Reagent (Life Technologies) per manufacturer’s instructions. First-strand cDNA was synthesized from 1 μg of total RNA using iScript Reverse Transcription Supermix for RT-PCR (Bio-Rad Laboratories) according to the manufacturer’s instructions. Individual 20 μl SYBR Green real-time PCR reactions consisted of 1 μl of diluted cDNA, 10 μl of SYBR Green Supermix (Bio-Rad), and 1 μl of each 5 μM forward and reverse primers. For standardization of quantification, 18S was amplified simultaneously. The PCR was carried out on 96-well plates on a CFX Connect Real time System (Bio-Rad), using a three-stage program provided by the manufacturer: 95 °C for 3 min, 40 cycles of 95 °C for 10 s and 60 °C for 30 s. Gene-specific primers are tabulated in Table S1.

### Quantification and Statistical Analysis

Statistical analyses were performed using Graphpad PRISM. Unless indicated, all results shown as mean ± SD of cellular triplicates obtained from one representative experiment as specified in each figure legend. P values were calculated using a Student’s two-tailed *t* test, One-way ANOVA w/ Dunnet’s method for multiple comparisons, or Two-way ANOVA w/ Tukey’s method for multiple comparisons; *, P value < 0.05; **, P value < 0.01; ***, P value < 0.001, ****, P value < 0.0001. Unless indicated, all normalization and statistical tests compared to wildtype cells.

**Table 1.** Key Resources Table.

| REAGENT or RESOURCE | SOURCE | IDENTIFIER |
| --- | --- | --- |
| Bacterial and virus strains |  |  |
| One Shot Stbl3 E. Coli | ThermoFisher Scientific | Cat#C7373 |
| Library Efficiency DH5α Competent Cells | ThermoFisher Scientific | Cat#18263012 |
| Chemicals, peptides, and recombinant proteins |  |  |
| [U-<sup>13</sup>C<sub>6</sub>] Glucose | Cambridge Isotope Labs | Cat#CLM-1396 |
| [U-<sup>13</sup>C<sub>5</sub>] Glutamine | Cambridge Isotope Labs | Cat#CLM-1822 |
| [U-<sup>13</sup>C<sub>2</sub>] Acetate | Cambridge Isotope Labs |  |
| [3-<sup>13</sup>C<sub>1</sub>] Pyruvate | Cambridge Isotope Labs |  |
| [2,4-<sup>13</sup>C<sub>6</sub>] Citrate | Cambridge Isotope Labs | Cat#CLM-148 |
| [U-<sup>13</sup>C<sub>8</sub>] Octanoate | Cambridge Isotope Labs | Cat#CLM-9617 |
| MTBSTFA + 1% TBDMSCI | Regis Technologies | Cat#1-270143-200 |
| BsmBI | New England Biolabs | Cat#R0580L |
| Critical commercial assays |  |  |
| Lipofectamine 3000 | ThermoFisher Scientific | Cat#L3000008 |
| KAPA HiFi HotStart Ready Mix | Kapa Biosystems | Cat#KK2602 |
| iTaq Universal SYBR Green | Biorad | Cat#1725121 |
| iScript Reverse Transcription Supermix | Biorad | Cat#1708840 |
| RNeasy Mini Kit | QIAGEN | Cat#74104 |
| Zero Blunt TOPO PCR Cloning Kit | ThermoFisher Scientific | Cat#450245 |
| prepGEM gold buffer | Zygem |  |
| Experimental models: cell lines |  |  |
| HepG2 | ATCC | Cat#HB-8065 |
| A549 | ATCC | Cat#CCL-185 |
| HAP1 | Horizon Discovery | Cat#C631 |
| 634T | Svensson et. al. (2016) | NA |
| HEK293FT | ThermoFisher Scientific | Cat#R70007 |
| Oligonucleotides |  |  |
| See Table S1 |  |  |
| Recombinant DNA |  |  |
| LentiCRISPR v2 plasmid | Addgene | Cat#52961 |
| pCMVR8.2 | Addgene | Cat#12263 |
| pMD2.G | Addgene | Cat#12259 |
| Software and algorithms |  |  |
| MATLAB 2016b | MathWorks | https://www.mathworks.com/products/matlab.html |
| Escher-Trace | Kumar et al. (2020) | https://escher-trace.github.io/ |
| GraphPad Prism 7.00 | GraphPad | https://www.graphpad.com/scientific-software/prism/ |
| INCA | Young et al. (2014) | https://mfa.vueinnovations.com/ |
| CRISPOR | Concordet et al. (2018) | http://crispor.tefor.net/ |
| NCBI BLAST | Agarwala et al. (2018) | https://blast.ncbi.nlm.nih.gov/ |

## Acknowledgments

We thank all members of the Metallo laboratory for support and helpful discussions. ACLY and ACSS2 cellular reagents were provided by Nimbus Therapeutics. Figure panels were made with Biorender.

## Funding

US National Institutes of Health (NIH) grants R35CA220538 and P01CA120964 (to R.J.S.) and R01CA234245 (to C.M.M.). A.K. was supported in part by U54CA132379, and H.G. was supported in part by T32DK007541.

## Author contributions

A.K., R.K., R.U.S. and C.M.M. conceived and designed the study. A.K. and R.K. performed experiments. R.K. generated the *PEX5*-KO cell lines. H.G. and R.J.S. provided 643T cell lines and advised on studies. G.H.M. and C.R.G. performed LC-MS/MS lipidomic analytics. T.C. performed the ferroptosis assay. A.K., R.K. and C.M.M wrote the paper with input from all authors.

## Competing interests

Robert Svensson is an employee of Nimbus Therapeutics. Christian Metallo is a scientific advisor for Faeth Therapeutics.

## Data and materials availability

This study did not generate new unique reagents. The datasets supporting the current study are available from the corresponding author upon request.

## Notes

### Competing Interest Statement

The authors have declared no competing interest.

### Summary of Updates

reverting to initial submission as requested by journal

