## Supplemental material for "Inter-organelle crosstalk supports acetyl-coenzyme A homeostasis and lipogenesis under metabolic stress"

**This PDF file includes:**

Figs. S1 to S6
Tables S1
Tables S2

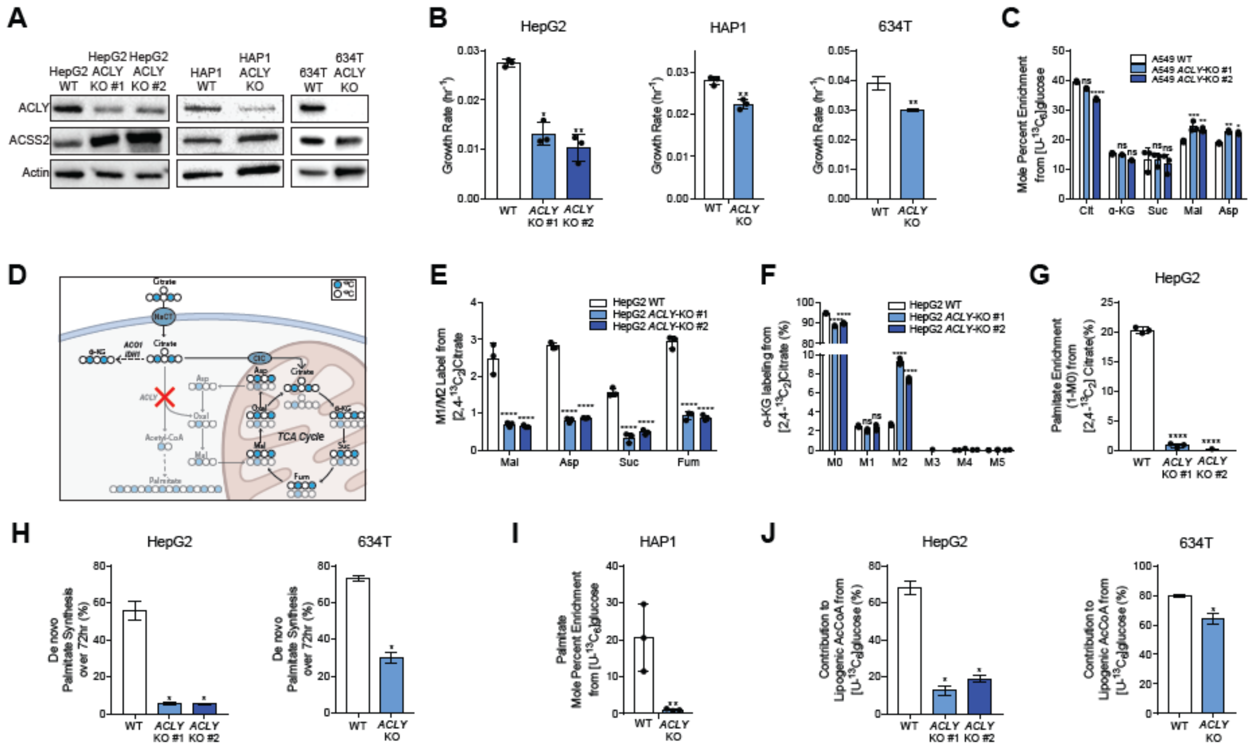


**Fig. S1.**

**Supplementary Figure 1. ACLY-KO rewires central carbon metabolism and leads to a reduction of palmitate synthesis in cancer cells, related to Figure 1.**
(A) Western blots of ACLY, ACSS2, and actin in WT and ACLY-KO HepG2, HAP1, and 634T cells.
(B) Growth rates of WT and ACLY-KO HepG2, HAP1, and 634T cells grown in high glucose DMEM +10% FBS for 4 days.
(C) Mole percent enrichment of TCA intermediates from [U-13C6]glucose in A549 ACLY-KO cells cultured in high glucose DMEM +10% dFBS for 72 hours (n=3).
(D) Schematic of extracellular [2,4-13C6]citrate catabolism. Filled circles are 13C, empty circles are 12C. Faded metabolite labeling and arrows are reduced by ACLY knockout.
(E-G) Ratio of the relative abundance of M1/M2 TCA intermediates from [2,4-13C2]citrate (E), Mass isotopomer distribution of α-KG from [2,4-13C2]citrate (F), Enrichment (1-M0) of palmitate from [2,4-13C2]citrate (G) in HepG2 WT and ACLY-KO cells cultured in high glucose DMEM +10% dFBS + 500 µM citrate for 72 hours (n=3).
(H) De novo synthesis of palmitate in WT and ACLY-KO HepG2 (left) and 634T (right) cells cultured in high glucose DMEM +10% dFBS (n=3).
(I) Mole percent enrichment of palmitate from [U-13C6] glucose in HAP1 WT and ACLY-KO cells cultured in high glucose DMEM +10%dFBS (n=3).
(J) Percent of lipogenic acetyl-CoA contributed by [U-13C6]glucose based on ISA modeling in WT and ACLY-KO HepG2 (left) and 634T (right) cells cultured in high glucose DMEM +10% dFBS (n=3).
In (B,C,E-G, and I) data are plotted as mean ± SD. Statistical significance is relative to WT as determined by One-way ANOVA w/Dunnet's method for multiple comparisons with *, P value < 0.05; **, P value < 0.01; ***, P value < 0.001, ****, P value < 0.0001. In (H, J) *data are plotted as mean estimated flux +/- 95% confidence intervals generated via parameter continuation in INCA. Unless indicated, all data represent biological triplicates. Data shown are from one of at least two separate experiments.

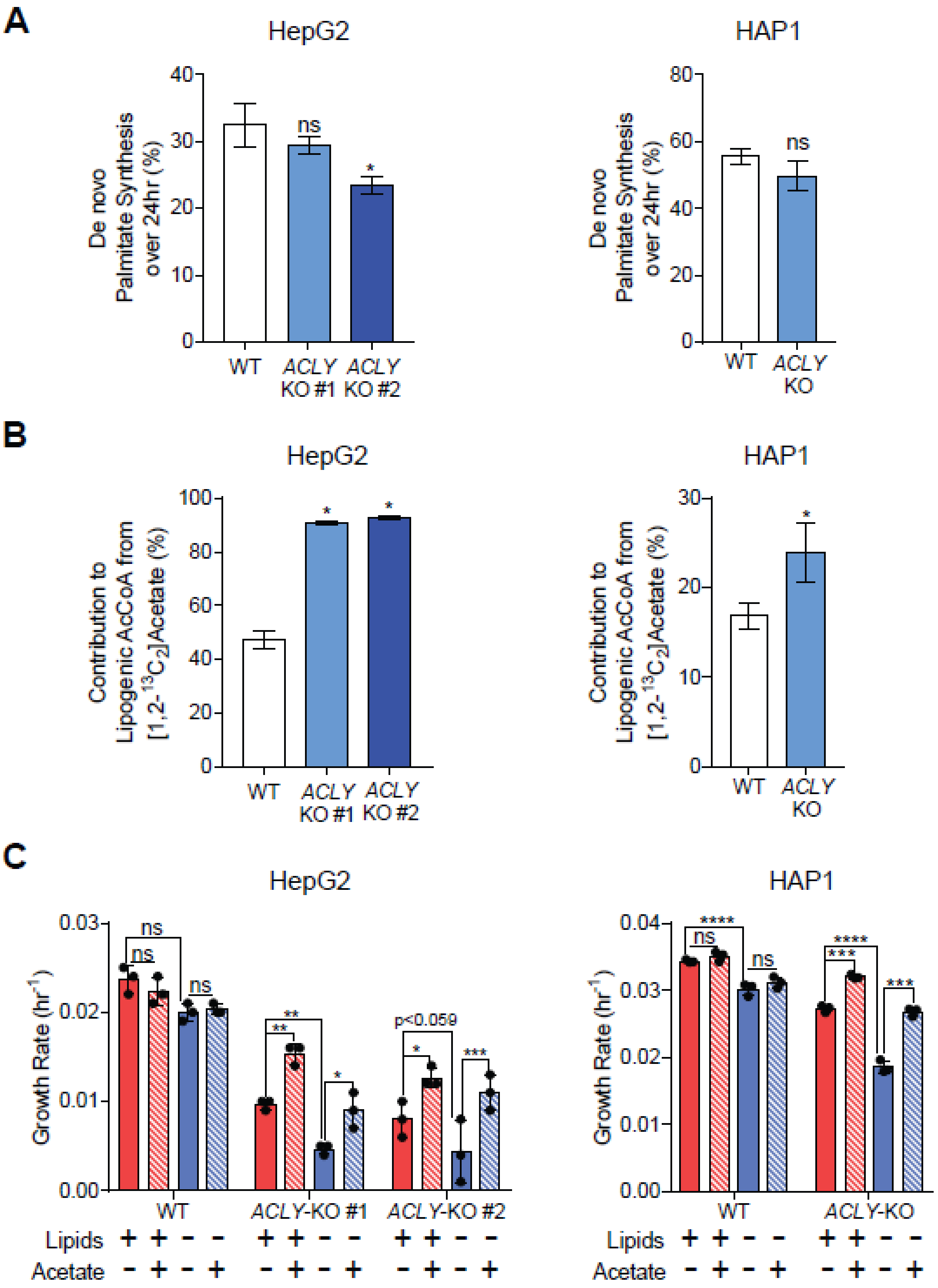
A
HepG2
De novo Palmitate Synthesis over 24hr (%)
WT
ACLY KO #1
ACLY KO #2
ns
*
HAP1
De novo Palmitate Synthesis over 24hr (%)
WT
ACLY KO
ns
B
HepG2
Contribution to Lipogenic AcCoA from [1,2-13C2]Acetate (%)
WT
ACLY KO #1
ACLY KO #2
HAP1
Contribution to Lipogenic AcCoA from [1,2-13C2]Acetate (%)
WT
ACLY KO
C
HepG2
Growth Rate (hr-1)
WT
ACLY-KO #1
ACLY-KO #2
p<0.059
Lipids + + - -
Acetate - + - +
HAP1
Growth Rate (hr-1)
WT
ACLY-KO
Lipids + + - -
Acetate - + - +

**Fig. S2.**

**Supplementary Figure 2. ACLY-KO growth and fatty acid synthesis is rescued with addition of extracellular acetate, related to Figure 2.**
(A) De novo synthesis of palmitate in HepG2 (left) and HAP1 (right) WT and ACLY-KO cells cells cultured in high glucose DMEM +10% dFBS + 1 mM acetate for 24 hours (n=3).
(B) Percent of lipogenic acetyl-CoA contributed by [1,2-13C2]acetate based on ISA modeling in HepG2 (left) and HAP1 (right) WT and ACLY-KO cells cultured in high glucose DMEM +10% dFBS + 1 mM acetate for 24 hours (n=3).
(C) Growth rates of HepG2 (left) and HAP1 (right) and WT and ACLY-KO cells grown in high glucose DMEM +10% dFBS or delipidated dFBS +/- 1 mM acetate for 4 days (n=3).
In (A, B) *data are plotted as mean estimated flux +/- 95% confidence intervals generated via parameter continuation in INCA. In
(C) data are plotted as mean ± SD. Statistical significance is determined by Two-way ANOVA w/ Tukey's method for multiple comparisons with *, P value < 0.05; **, P value < 0.01; ***, P value < 0.001, ****, P value < 0.0001.
Unless indicated, all data represent biological triplicates. Data shown are from one of at least two separate experiments.

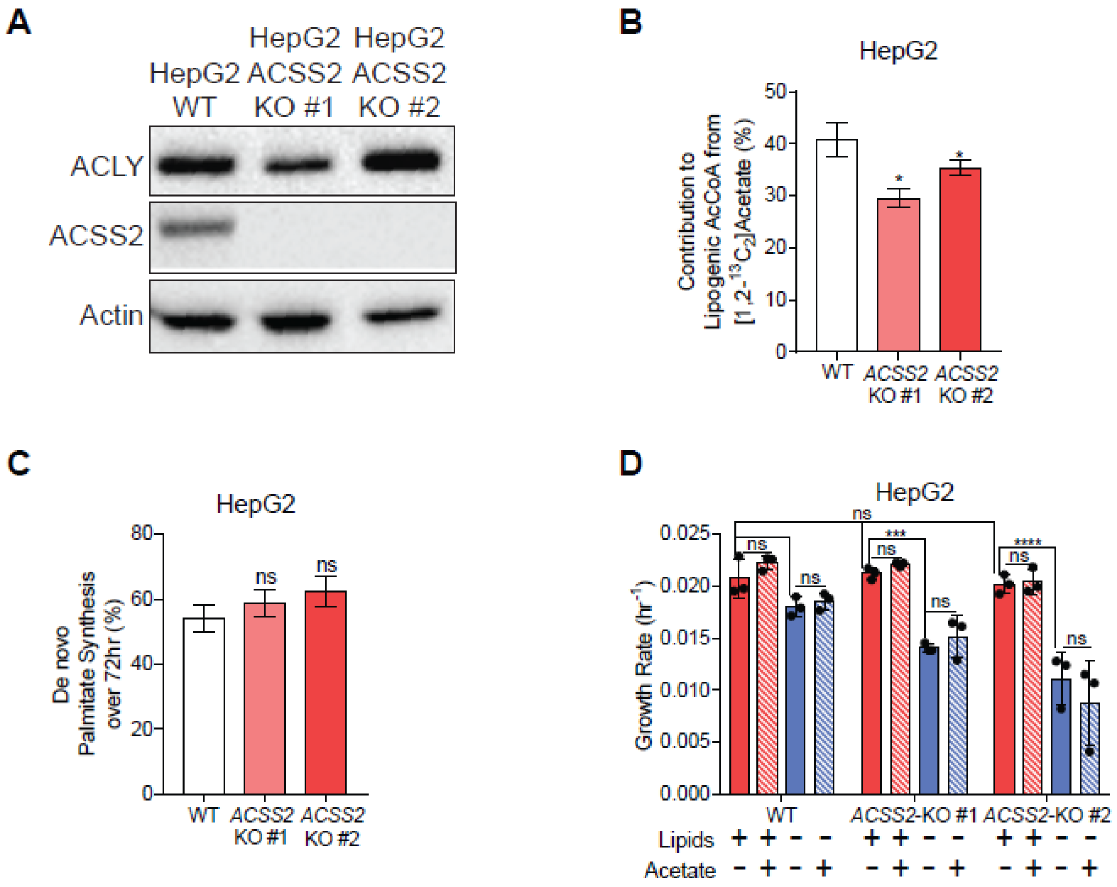


### Fig. S3.

**Supplementary Figure 3. ACSS2-KO attenuates catabolism of exogenous acetate with minimal effect on glucose catabolism, related to Figure 3.**
(A) Western Blots of ACLY, ACSS2, and actin in HepG2 WT and ACSS2-KO cells.
(B) Percent of lipogenic acetyl-CoA contributed by [1,2-13C2] acetate based on ISA modeling in HepG2 ACSS2-KO cells cultured in high glucose DMEM +10% dFBS + 1 mM acetate for 24 hours (n=3).
(C) De novo synthesis of palmitate in HepG2 ACSS2-KO cells cultured in high glucose DMEM +10% dFBS for 72 hours (n=3).
(D) Growth rates of HepG2 WT and ACSS2-KO cells grown in high glucose DMEM +10% dFBS or delipidated dFBS +/- 1 mM acetate for 4 days (n=3).

In (B, C) *data are plotted as mean estimated flux +/- 95% confidence intervals generated via parameter continuation in INCA. In (D) data are plotted as mean ± SD. Statistical significance is determined by Two-way ANOVA w/ Tukey's method for multiple comparisons D) with *, P value < 0.05; **, P value < 0.01; ***, P value < 0.001, ****, P value < 0.0001. Unless indicated, all data represent biological triplicates. Data shown are from one of at least two separate experiments.

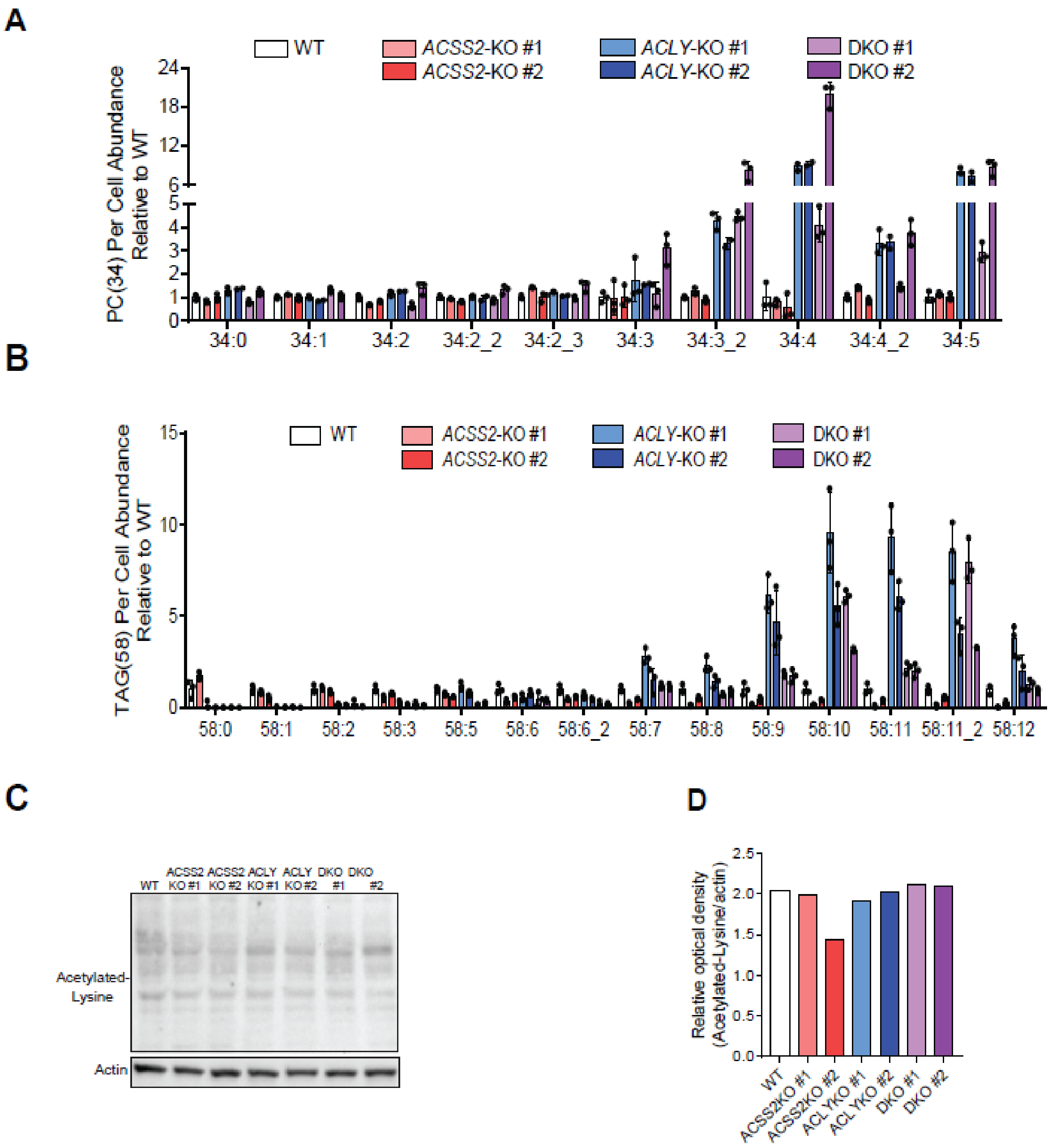


**Fig. S4.**

**Supplementary Figure 4. ACLY/ACSS2- cells are reliant on extracellular lipids, with minimal change in protein acetylation, related to Figure 4.**
(A) PC(34) per cell abundance of A549 WT, ACSS2-KO, ACLY-KO, ACLY/ACSS2-DKO cells cultured in high glucose DMEM +10% FBS for 2 days (n=3).
(B) TAG(58) per cell abundance of A549 WT, ACSS2-KO, ACLY-KO, ACLY/ACSS2-DKO cells cultured in high glucose DMEM +10% FBS for 2 days (n=3). Data are plotted as mean ± 95% confidence interval (CI). Unless indicated, all data represent biological triplicates.
(C) Western Blots of acetylated-lysine and actin in A549 WT, ACSS2-KO, ACLY-KO, ACLY/ACSS2 DKO whole cell lysates.

(D) Relative optical density of acetylated-lysine/actin in A549 WT, ACSS2-KO, ACLY-KO, ACLY/ACSS2-DKO whole cell lysates (n=1).

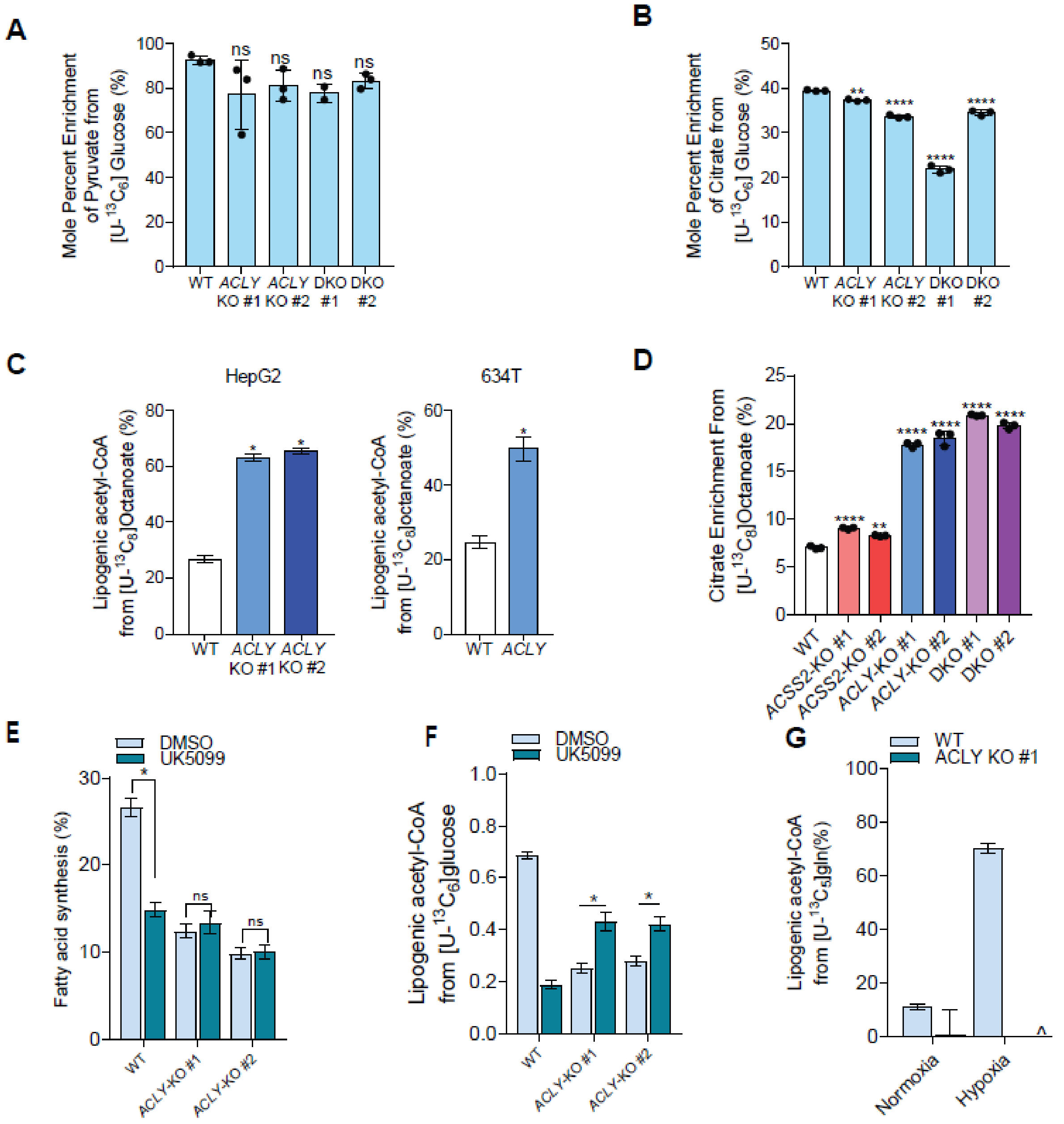


**Fig. S5.**

**Supplementary Figure 5. Disruption of canonical acetyl-CoA synthesis induces alternative synthesis pathways, related to Figure 5.**
(A-B) Mole percent enrichment of pyruvate (A) and citrate (B) from [U-13C6] glucose in A549 WT, ACLY-KO, ACLY/ACSS2-DKO cells cultured in high glucose DMEM +10% dFBS for 72 hours (n=3).
(C) Percent of lipogenic acetyl-CoA contributed by [U-13C8] octanoate based on ISA modeling in WT and ACLY-KO HepG2 (left) and 634T (right) cells cultured in high glucose DMEM +10% dFBS + 500 µM octanoate for 24 hours. (n=3).
(D) Mole percent enrichment of citrate from [U-13C8] octanoate in A549 WT, ACSS2-KO, ACLY-KO, ACLY/ACSS2-DKO cells cultured in high glucose DMEM +10% dFBS + 500 µM octanoate for 24 hours (n=3).

(E-F) De novo synthesis of palmitate (E), and Percent of lipogenic acetyl-CoA contributed by [U-13C6] glucose based on ISA modeling (F) in A549 WT, and ACLYKO cells cultured in DMEM +10% dFBS and treated with 20uM MPC for 48 hours. (n=3)

(G)Percent of lipogenic acetyl-CoA contributed by [U-13C5] glutamine based on ISA modeling in A549 WT, and ACLYKOcells cultured in high glucose DMEM +10% dFBS + 4mM [U-13C5] glutamine for 72 hours under hypoxia or normoxia. (n=3)

In (A,B,D) Data are plotted as mean ± SD. Statistical significance relative to WT as determined by One-way ANOVA w/ Dunnet's method for multiple comparisons with *, P value < 0.05; **, P value < 0.01; ***, P value < 0.001, ****, P value < 0.0001. In (C,,E-G) *data are plotted as mean estimated flux +/- 95% confidence intervals generated via parameter continuation in INCA. Unless indicated, all data represent biological triplicates. Data shown are from one of at least two separate experiments.

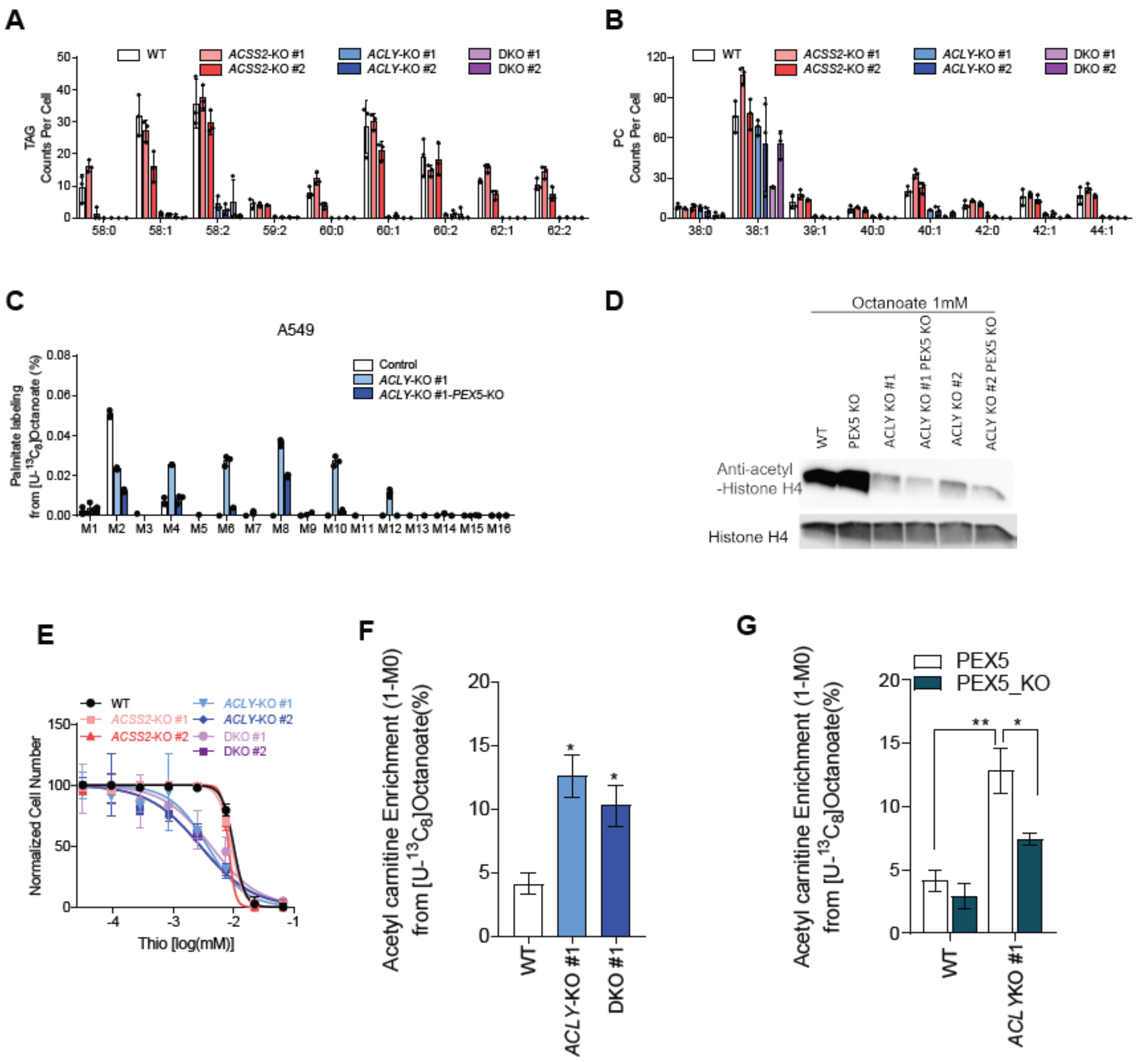


**Fig. S6.**

**Supplementary Figure 6. Peroxisomal β-oxidation becomes a major source of lipogenic acetyl-CoA with ACLYKO and ACLY/ACSS2 DKO, related to Figure 6.**
(A) Saturated, mono- and di-unsaturated TAG per cell abundances of A549 WT, ACSS2-KO, ACLY-KO, ACLY/ACSS2-DKO cells cultured
in high glucose DMEM +10% FBS 2 days (n=3).
(B) Saturated and monounsaturated PC per cell abundances of A549 WT, ACSS2-KO, ACLY-KO, ACLY/ACSS2-DKO cells cultured in high
glucose DMEM +10% FBS 2 days (n=3).
(C) Mass isotopomer distribution of palmitate from [U-13C8] octanoate in A549 WT, ACLY-KO and ACLY/PEX5-DKO cells cultured in high glucose DMEM +10% dFBS + 500 µM octanoate for 48 hours (n=3).
(D) Western Blots of Anti-acetyl-Histone H4 and Histone H4 in A549 WT, PEX5 KO, ACLY KO and ACLY/PEX5-DKO cells.
(E) Dose response of A549 WT, ACSS2-KO, ACLY-KO, ACLY/ACSS2-DKO cells cultured in high glucose DMEM +10% dFBS to Thioridazine over 4 days, x-axis is log10 (n=3).
(F)Enrichment (1-M0) of acetyl carnitine from [U-13C8] octanoate in A549 WT, ACLY-KO, ACLY/ACSS2-DKO cells cultured in high glucose DMEM +10% dFBS for 48 hours (n=3)
(G) Enrichment (1-M0) of acetyl carnitine from [U-13C8] octanoate in A549 WT, ACLY-KO, ACLY/PEX5-KO cells cultured in high glucose DMEM +10% dFBS for 48 hours (n=3).
Unless indicated, all data represent biological triplicates. Data shown are from one of at least two separate experiment.

### Table S1.

Orbitrap high-resolution (QE) mass spectrometry lipid analysis.

| Lipid species | Precursor ion | Product ion |
|---|---|---|
| ***Triacylglycerol (TAG)*** | | |
| TAG 44:0 | 768.7083 | 495.4 |
| TAG 44:1 | 766.6931 | 495.4 |
| TAG 44:2 | 764.6774 | 465.4 |
| TAG 46:0 | 796.7404 | 523.5 |
| TAG 46:1 | 794.724 | 521.5 |
| TAG 46:2 | 792.7076 | 521.5 |
| TAG 47:0 | 810.7551 | 537.5 |
| TAG 47:1 | 808.7396 | 535.5 |
| TAG 47:2 | 806.7241 | 535.5 |
| TAG 48:0 | 824.7709 | 551.5 |
| TAG 48:1 | 822.7551 | 549.5 |
| TAG 48:2 | 820.7398 | 549.5 |
| TAG 48:3 | 818.7241 | 547.5 |
| TAG 49:0 | 838.7861 | 565.5 |
| TAG 49:1 | 836.7688 | 563.5 |
| TAG 49:2 | 834.756 | 563.5 |
| TAG 50:0 | 852.8017 | 579.5 |
| TAG 50:1 | 850.7866 | 577.5 |
| TAG 50:2 | 848.7709 | 575.5 |
| TAG 50:3 | 846.7551 | 575.5 |
| TAG 50:4 | 844.7398 | 577.5 |
| TAG 52:0 | 880.8356 | 607.5 |
| TAG 52:1 | 878.8177 | 605.5 |
| TAG 52:2 | 876.8022 | 577.5 |
| TAG 52:3 | 874.7866 | 575.5 |
| TAG 52:4 | 872.7714 | 551.5 |
| TAG 52:5 | 870.7556 | 549.5 |
| TAG 52:6 | 868.7409 | 523.5 |
| TAG 53:1 | 892.8328 | 591.5 |
| TAG 53:2 | 890.8173 | 591.5 |
| TAG 53:3 | 888.8032 | 589.5 |
| TAG 54:0 | 908.8629 | 607.5 |
| TAG 54:1 | 906.8494 | 633.6 |
| TAG 54:2 | 904.8335 | 605.5 |
| TAG 54:3 | 902.8179 | 603.5 |
| TAG 54:4 | 900.8021 | 601.5 |
| TAG 54:5 | 898.7863 | 551.5 |
| TAG 54:6 | 896.7711 | 551.5 |
| TAG 54:7 | 894.7564 | 549.5 |
| TAG 54:8 | 892.7405 | 547.5 |
| TAG 55:2 | 918.8492 | 619.6 |

| | | |
|---|---|---|
| TAG 55:3 | 916.8337 | 617.5 |
| TAG 55:5 | 912.8025 | 591.5 |
| TAG 55:6 | 910.7889 | 565.5 |
| TAG 55:7 | 908.7744 | 563.5 |
| TAG 56:0 | 936.8923 | 663.5 |
| TAG 56:1 | 934.8777 | 661.6 |
| TAG 56:2 | 932.8624 | 633.6 |
| TAG 56:3 | 930.8469 | 631.6 |
| TAG 56:4 | 928.8312 | 629.5 |
| TAG 56:5 | 926.8129 | 627.5 |
| TAG 56:6 | 924.7978 | 577.5 |
| TAG 56:7 | 922.7868 | 577.5 |
| TAG 56:8 | 920.7715 | 575.5 |
| TAG 56:9 | 918.7554 | 573.5 |
| TAG 57:7 | 936.8024 | 591.5 |
| TAG 57:8 | 934.7869 | 589.5 |
| TAG 58:0 | 964.9256 | 691.7 |
| TAG 58:1 | 962.9064 | 663.6 |
| TAG 58:2 | 960.8938 | 661.6 |
| TAG 58:3 | 958.8768 | 659.6 |
| TAG 58:5 | 954.8446 | 605.5 |
| TAG 58:6 | 952.8306 | 605.5 |
| TAG 58:7 | 950.8155 | 605.5 |
| TAG 58:8 | 948.8008 | 603.5 |
| TAG 58:9 | 946.7828 | 601.5 |
| TAG 58:10 | 944.7669 | 599.5 |
| TAG 58:11 | 942.7509 | 597.5 |
| TAG 58:12 | 940.7402 | 595.5 |
| TAG 59:2 | 974.9122 | 577.5 |
| TAG 60:0 | 992.9558 | 691.7 |
| TAG 60:1 | 990.9384 | 605.5 |
| TAG 60:2 | 988.9228 | 689.6 |
| TAG 60:3 | 986.9088 | 603.5 |
| TAG 60:6 | 980.8616 | 605.5 |
| TAG 60:7 | 978.8457 | 605.5 |
| TAG 60:8 | 976.8324 | 631.5 |
| TAG 60:9 | 974.8166 | 629.5 |
| TAG 60:10 | 972.7974 | 627.5 |
| TAG 60:11 | 970.7842 | 625.5 |
| TAG 62:1 | 1018.971 | 719.7 |
| TAG 62:2 | 1016.956 | 717.7 |
| TAG 62:3 | 1014.942 | 715.7 |
| TAG 62:6 | 1008.892 | 605.5 |
| TAG 62:7 | 1006.877 | 706.6 |

#### *Phosphatidylcholine (PC)*

| | | |
|---|---|---|
| PC 33:1 | 746.568 | 184.0734 |

| | | |
|---|---|---|
| PC 33:0 | 748.5834 | 184.0734 |
| PC 34:0 | 762.5991 | 184.0734 |
| PC 34:1 | 760.5837 | 184.0734 |
| PC 34:2 | 758.5677 | 184.0734 |
| PC 34:3 | 756.5516 | 184.0734 |
| PC 36:0 | 790.6306 | 184.0734 |
| PC 36:1 | 788.6147 | 184.0734 |
| PC 36:2 | 786.5991 | 184.0734 |
| PC 36:3 | 784.5825 | 184.0734 |
| PC 36:4 | 782.5677 | 184.0734 |
| PC 36:5 | 780.5502 | 184.0734 |
| PC 36:6 | 778.5353 | 184.0734 |
| PC 38:1 | 816.6465 | 184.0734 |
| PC 38:2 | 814.6306 | 184.0734 |
| PC 38:3 | 812.6135 | 184.0734 |
| PC 38:4 | 810.5991 | 184.0734 |
| PC 38:5 | 808.5831 | 184.0734 |
| PC 38:6 | 806.5675 | 184.0734 |
| PC 38:7 | 804.5498 | 184.0734 |
| PC 40:0 | 846.6929 | 184.0734 |
| PC 40:1 | 844.6782 | 184.0734 |
| PC 42:0 | 874.7244 | 184.0734 |
| PC 42:1 | 872.7089 | 184.0734 |
| PC 44:0 | 902.7556 | 184.0734 |
| PC 44:1 | 900.7404 | 184.0734 |

**Table S2.**

| Isotopomer spectral analysis | Normalized flux | 95% confidence interval | |
|---|---|---|---|
| Pathway/Reaction | | Lower bound | Upper bound |
| Palmitate ISA | | | |
| Ac.l -> Ac (Lipogenic acetyl CoA) | 0.6369 | 0.5675 | 0.6973 |
| Ac.d -> Ac | 0.3631 | 0.3027 | 0.4325 |
| Ac + Ac + Ac + Ac + Ac + Ac + Ac + Ac -> Palm.s | 0.125 | 0.125 | 0.125 |
| 0*Palm.s -> Palm (Fatty acid synthesis) | 0.2562 | 0.2194 | 0.2902 |
| Palm.d -> Palm | 0.7438 | 0.7098 | 0.7806 |
| Palm -> Palm.m | 1 | 1 | 1 |
| Palm.s -> Palm.f | 0.125 | 0.125 | 0.125 |

270 m/z fragment ion of palmitate methylester was used for isotopomer spectral analysis
